# Stigma Receptivity is controlled by Functionally Redundant MAPK Pathway Components in Arabidopsis

**DOI:** 10.1101/2020.03.09.983767

**Authors:** Muhammad Jamshed, Subramanian Sankaranarayanan, Kumar Abhinandan, Marcus A. Samuel

**Affiliations:** University of Calgary, BI 392, Department of Biological Sciences, 2500 University Dr. NW. Calgary, Alberta T2N 1N4, Canada.; Senior Scientist, Frontier Agri-Science Inc, 98 Ontario Street. Port Hope, Ontario, L1A 2V2, Canada.; Department of Botany and Plant Pathology, Purdue University, West Lafayette, 47907, Indiana, USA.

## Abstract

In angiosperms, the process of pollination relies on species-specific interaction and signaling between the male (pollen) and female (pistil) counterparts where the interplay between several pollen and stigma proteins decides the fate of the pollen. In Brassicaceae, the dry stigmatic papillary cells control pollen germination by releasing resources only to compatible pollen thereby allowing pollen to hydrate and germinate. Despite the identification of a number of stigmatic proteins that facilitate pollination responses, the signaling mechanisms that regulate functions of these proteins have remained unknown. Here we show that, in Arabidopsis, an extremely functionally redundant mitogen-activated protein kinase (MAPK) cascade is required for maintaining stigma receptivity to accept compatible pollen. Our genetic analyses demonstrate that in stigmas, five MAPK kinases (MKKs), MKK1/2/3/7/9 are required to transmit upstream signals to two MPKs, MPK3/4, to mediate compatible pollination. Compromised functions of these five *MKKs* in the quintuple mutant (*mkk1/2/3RNAi/mkk7/9*) phenocopied pollination defects observed in the *mpk4RNAi/mpk3* double mutant. We further show that this MAPK nexus converges on Exo70A1, a previously identified stigmatic compatibility factor essential for pollination. Given that pollination is the crucial initial step during plant reproduction, understanding the mechanisms that govern successful pollination could lead to development of strategies to improve crop yield.

## Introduction

Pollination in angiosperms is a highly coordinated process which relies on proper cell-cell communication and signaling between the pollen grain and the stigmatic papillae. In Brassicaceae, the female part of the flower regulates pollination through stigma components called compatibility factors, which are induced as a result of interaction with a suitable mate (pollen) (1). These compatibility factors are proposed to facilitate transport of water and nutrition for pollen hydration, germination and pollen tube growth (2–4)1. However, the signaling mechanism behind how these factors are regulated has remained largely unknown. Interestingly, during compatible interactions, calcium spikes and constitutive ROS accumulation are observed in the stigmas, which most likely can trigger other compatibility factors necessary for pollen growth and germination (5–7). There is also precedence for both these second messengers as strong activators of Mitogen-Activated Protein Kinases (MAPK) (8, 9).

MAPK cascade is a highly conserved signaling network in eukaryotes which is involved in signal transduction of inputs from various environmental and developmental stimuli (10). The MAPK phosphorylation cascade operates in a hierarchical fashion with activated MAPKKK(s) phosphorylating and activating MAPKK(s), which in turn activate MAPK(s) (11). In Arabidopsis, there are 10 MAPKK(s) (also known as MKKs) and 20 MAPK(s) (commonly known as MPKs), which control numerous cellular responses by regulating different enzymes and transcription factors (11). This allows for a staggering number of possible combinations of MKK-MPK pairs with the addition of possible functional redundancy amongst these proteins. During plant reproduction, various MAPK cascades have been shown to control floral architecture, anther development, ovule development and pollen tube guidance (12–16). Despite this, the role of MAPKs during early stages of pollination has not been explored.

Interestingly, mammalian orthologs of TEY class of plant MAPKs, ERK1/2, have been shown to phosphorylate Exo70, a member of exocyst complex, which is involved in exocytosis through tethering, docking and fusion of secretory vesicles to plasma membrane(17). This phosphorylation was proposed to promote post-Golgi vesicle transport to plasma membrane to regulate exocytosis along with other cellular processes in response to growth factor signaling (17). In Arabidopsis and canola, Exo70A1 plays an essential role during pollination likely through regulating delivery of vesicles to the plasma membrane required for pollen germination (18).

Given that both calcium and ROS strong activators of MAPK signaling (8, 9), we predicted that MAPKs will be constitutively active in the stigmatic papillary cells and could play a role during pollination responses. In agreement with our hypothesis, treatment of flowers with a MEK1/2 (MAPKK) inhibitor, U0126 (19) previously successfully used to inhibit activation of plant MAPKs (9, 20), caused a significant reduction in compatible pollen attachment and germination (Supplemental Fig. S1A, B) suggesting the involvement of the MAPK family in regulating pollination. Our genetic analysis of the MAPK pathway components in Arabidopsis revealed a functionally redundant MKK/MPK network that converges on Exo70A1 to mediate pollination responses.

## Results

### MAPKs are constitutively active in mature stigmas

While the U0126 inhibitor study provided preliminary evidence for role for MAPKs in pollination, the challenge was to identify which of the 20 MPKs were involved. We took a biochemical approach using in-gel kinase assays to identify MPKs that are active in the stigmas from stage 12 flowers which undergo anthesis and are primed to accept pollen. When proteins from unpollinated stigmas of stage 12 Arabidopsis Col-0 flowers were subjected to in-gel kinase assays, we identified three kinases that were active in mature stigmas (Fig. 1A). The masses of these three kinases and previous evidence with in-gel kinase assays using other Arabidopsis tissue types, suggested to us that these three kinases could be MPK3, MPK4 and MPK6 (11, 21). Two of these kinases were validated as MPK6 and MPK3 through using proteins from stigmas of homozygous SALK transfer (T)-DNA mutant lines of *mpk6-2* (SALK_073907) and *mpk3-1* (SALK_151594) where the respective bands were absent (Fig. 1A). Since Arabidopsis *mpk4* mutant is a dwarf plant with severe floral defects (22), we suppressed *MPK4* in the stigmas through an RNAi approach using the stigma-specific *SLR1* promoter. Our in-gel kinase assays with stigmatic proteins from *SLR1::MPK4Ri* plants validated the identity of the third kinase as MPK4 (Fig. 1B).

**Fig. 1.**
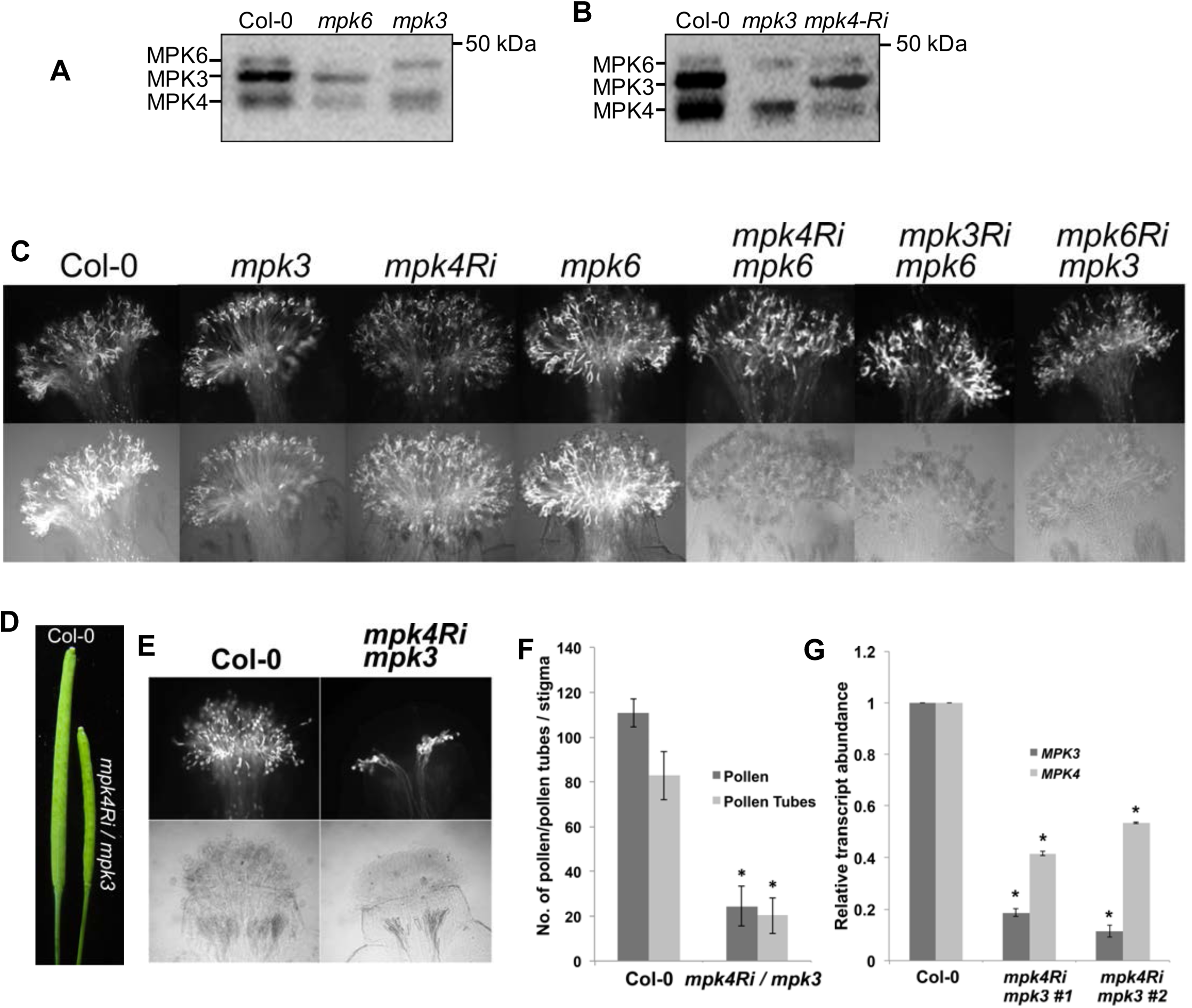
Loss of MPK3 and MPK4 results in reduced compatibility. **(A)** In-gel kinase assays showing activity of MPK3 and MPK6 in Col-0, *mpk3* and *mpk6* mutant in stage 12 stigmas. **(B)** In-gel kinase assay showing activity of MPK4 and MPK3 in Col-0, *mpk3*, *mpk4 RNAi* stage 12 stigmas. **(C)** Aniline blue assays of stigmas from various MAPK single and digenic mutants showing pollen and pollen tubes following self-pollination. All stigmas were allowed to naturally pollinate and collected four hours after flower opening. All RNAi constructs were placed under stigma specific *SLR1* promoter. **(D)** Siliques of Col-0 and *mpk4Ri/mpk3* mutant showing reduced seed set. **(E)** Aniline blue assays of pollinated stigmas from Col-0 and *mpk4Ri/mpk3* mutant. **(F)** Average number of pollen attached, and pollen tubes germinated in Col-0 and *mpk4Ri/mpk3* mutant (n>5). Error bars represents ± SE. Asterisk represents values significantly different from Col-o at P<0.05. **(G)** q-PCR analysis showing relative transcript abundance of *MPK3* and *MPK4* in stigmas from stage 12 flowers of Col-0 and two independent *mpk4Ri/mpk3* mutant lines (n=3). Error bars indicate ± SE. Asterisks indicate significant differences (*P<0.05).

### MPK3 and MPK4 play a functionally redundant role during pollination

Given the complexity of the MAPK pathway and the essential nature of MPK3/4/6 for development and cell survival (21, 22), we took a genetic approach combined with stigma-specific suppression to address the role of the MAPK cascade during pollination. Since single T-DNA mutants of *mpk6, mpk3-1 or mpk3-DG* (23), and *SLR::MPK4Ri* lacked any pollination phenotypes, we created all possible combinations of double mutants and the triple mutant (Fig. 1C, Supplemental Fig. S2A, Supplemental Table 1). Because of the highly sensitive nature of MAPKs to external stimuli including touch (21, 24) and in order to keep the perturbation to a minimum, we allowed newly opened flowers to naturally self-pollinate (synchronized with flower opening) and stigmas were collected to assess pollination responses. As can be observed from Fig. 1C, *mpk4Ri/mpk6, mpk3Ri/mpk6* and *mpk6Ri/mpk3* were found to be normal without any change in pollen acceptance (Supplemental Table. 1). However, one of the combinations of double mutants *mpk4Ri/mpk3 (mpk3-DG)* produced shorter siliques with almost 50 % reduction in seeds (Fig. 1D, Supplemental Fig. S2B). This mutant lacked any floral defects, produced homomorphic flowers, viable pollen and had efficient transfer of pollen to the stigma. (Supplemental Fig. S2C - F). Consistent with the low seed set, *mpk4Ri/mpk3* plants showed impaired stigma function when tested for pollination responses (Fig. 1E, F). After 4 h of pollination, *mpk4Ri/mpk3* stigmas displayed severely reduced pollen attachment and germination compared to wild-type Col-0 pistils (Fig. 1E, F). When stigmas from two independent *mpk4Ri/mpk3* lines were assessed for *MPK3* and *MPK4* expression, significantly reduced transcript levels of *MPK3* and *MPK4* (RNAi-suppressed) were observed relative to Col-0 (Fig. 1G). The triple mutant *mpk4Ri/mpk6Ri/mpk3* also mimicked the phenotype of *mpk4Ri/mpk3*, confirming the lack of a role for MPK6 during pollination (Supplemental Fig. S3A, B). Collectively, our observations indicate that both MPK3 and MPK4 are a functionally redundant pair of MAPKs required for mediating stigma receptivity.

### Five MKKs (MKK1/2/3/7/9) are required for pollination responses

We next investigated which of the ten upstream MKKs were necessary to transmit the signal to MPK3/4 to control pollination. For this, we used a targeted genetic approach to circumvent any lethality associated with loss of some of these kinases (10). Based on previous large-scale yeast two-hybrid and protein array data, eight MKKs (MKK1-6, 7&9) have the ability to interact or activate either MPK3 or MPK4 or both (25, 26). We did not include MKK6 in the screen since it is primarily known to be involved during cytokinesis (27). As shown in supplemental Fig. S4A, B; Fig. S5A-F, all possible combinations of these kinases were created followed by assessment of T1 and T2 generations for plants that phenocopied *mpk3/4* pollination phenotype (Supplemental Table 2). Of all the combinations tested, one of the quintuple mutants, *mkk1/2/3Ri/mkk7/9,* in which *MKK1/2/3* were suppressed in the stigmas of *mkk7/9* null mutant background, displayed pollination and fertility defects similar to *mpk3/4* (Fig. 2A-C, Supplemental Fig. S6A). This multi-locus mutant did not display any floral phenotypes and produced homomorphic flowers with normal anther dehiscence, viable pollen and unimpaired pollen transfer on to stigmas (Supplemental Fig. S6B-E). Loss or suppression of the respective *MKKs* was confirmed in mature stigmas of two independent quintuple mutant lines through qPCR (Fig. 2D). In *mkk1/2/3Ri/mkk7/9* line, suppression of *MKK3* was required to confer a phenotype identical to *mpk3/4* as the quadruple *mkk1/2Ri/mkk7/9* plants showed only moderate pollination defects, while the rest of the multi-locus mutants were fully fertile exhibiting no pollination anomalies (supplementary Fig. S5A-F; Fig. S4 A,B; Supplemental Table 2). This was not surprising, as MKK3 has been previously shown to activate MPK3 and MPK4 (26).

**Fig. 2.**
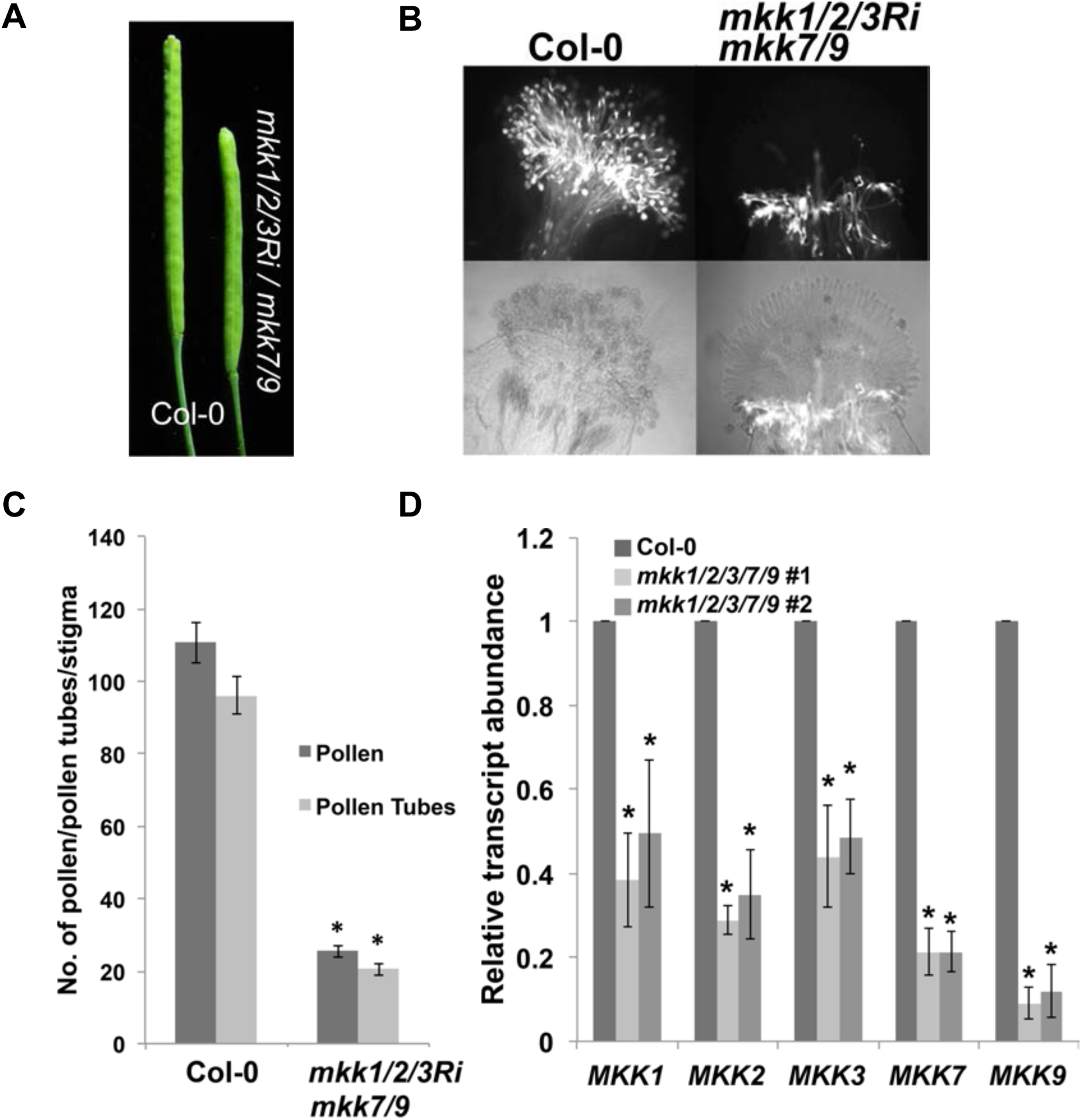
Five MKKs regulate compatible pollination. **(A)** Siliques of Col-0 and *mkk1/2/3Ri/mkk7/9* mutant showing reduced seed set. **(B)** Aniline blue assays of pollinated stigmas from Col-0 and *mkk1/2/3Ri/mkk7/9* mutant showing pollen and pollen tubes. **(C)** Average number of pollen attached, and pollen tubes germinated in Col-0 and *mkk1/2/3Ri/mkk7/9* mutant (n>5). Error bars shows ± SE. Asterisks indicate significant differences (*P<0.05). **(D)** q-PCR analysis showing relative transcript abundance of *MKK1,2,3,7* and *9* in stigmas from stage 12 flowers of Col-0 and two independent *mkk1/2/3Ri/mkk7/9* mutant lines (n=3). Error bars indicate ± SE. Asterisks indicate significant differences (*P<0.05).

### Activated MPK3 and MPK4 can phosphorylate the ‘SP’ motif of Exo70A1

Having established that MAPK signaling is required for compatible pollination to occur, we next focused on how MAPK signaling is integrated to regulate stigmatic factors required for pollen acceptance. One of the compatibility factors that has been shown to play a central role in stigma receptivity is Exo70A1, a member of the exocyst complex (18). Loss of *Exo70A1* resulted in inability of stigmas to support pollen hydration and germination (18). Protein sequence analysis of Exo70A1 revealed a single consensus MAPK phosphorylation (SP) motif with a docking domain upstream of the ‘SP’ motif (Supplementary Fig. S7). To test, if activated MPK4 can phosphorylate Serine 328 of the ‘SP’ motif of Exo70A1, full-length recombinant GST-Exo70A1 and GST-Exo70A1 (S328A) proteins were incubated with recombinant 6xHIS-MPK4 along with constitutively active GST-MKK1 (CAKK1) in the presence of radioactive γ ^32^P (Fig. 3A). Strong phosphorylation of GST-Exo70A1 was observed in the presence of 6xHIS-MPK4 and GST-CAKK1 and only marginal background phosphorylation was observed with GST-Exo70A1 (S328A) (Fig. 3A). To unequivocally demonstrate that Serine328 of Exo70A1 is phosphorylated by activated upstream MPK3/4, we obtained synthesized short peptides comprising of residues of 323 to 335 (AR***<u>S</u>***KR***<u>S</u>***PEKLFVL and AR***<u>S</u>***KR***<u>A</u>***PEKLFVL) with either an intact S328 residue or S328 mutated to an alanine and subjected them to in-vitro phosphorylation assays. These peptides were immobilized on PVDF membrane and subjected to in-vitro phosphorylation assays using activated 6xHIS-MPK4 (activated with GST-CAKK1) followed by immunoblotting with anti-phosphoserine antibodies (Fig. 3B). We observed that the synthetic peptide with an intact ‘SP’ motif was exclusively phosphorylated by activated MPK4 while the mutated S328A peptide was not detected by the phospho-serine antibodies (Fig. 3B). The results clearly demonstrate that MPK4 is able to specifically phosphorylate Serine328 of Exo70A1 (Fig. 3B). Similar ‘SP’-specific phosphorylation was also observed with activated MPK3 proteins (Fig. S8).

**Fig. 3.**
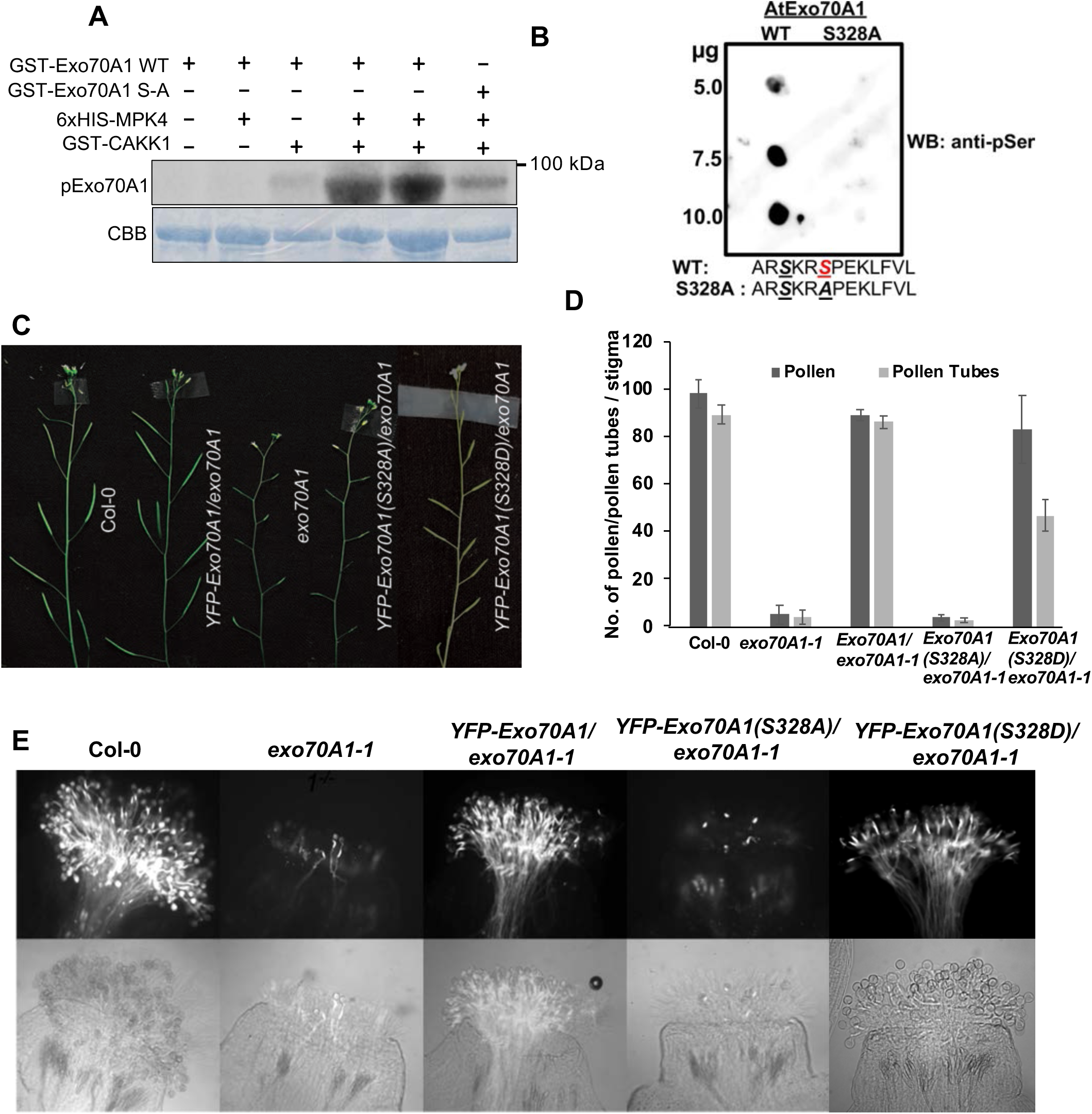
Phosphorylation of Exo70A1 is required for compatible pollination. **(A)** *In vitro* kinase assay showing phosphorylation of GST-Exo70A1 by 6xHIS-MPK4. Lanes 4 & 5 contain 5 and 10 μg of Exo70A1 respectively. **(B)** Dot blot showing the phosphorylation of WT peptide only blotted in various concentrations. Kinase reaction was run in the presence pre-activated 6xHIS-MPK4 with GST CAKK1 and blotted with anti-pSer antibody. **(C)** Inflorescence phenotype of Col-0, *35S::YFP-Exo70A1(WT)/exo70A1,exo70A1,35S::YFPExo70A1(S328A)/exo70A1* (Line# 2) and *35S::YFP-Exo70A1(S328D)/exo70A1* (Line# 41). **(D)** Average number of pollen attached, and pollen tubes germinated on stigmas of these lines. Error bars indicate ± SE (n>5). Asterisks indicate significant differences (*P<0.05). **(E)** Aniline blue images of pollinated stigmas from respective WT, mutant and complemented lines.

### Exo70A1 Serine 328 of ‘SP’ motif is required for successful pollination

In order to further characterize whether this MAPK-mediated phosphorylation of Serine328 of Exo70A1 is needed for its function in pollination, we used complementation assays to transform *exo70A1-1* with either *35S::YFP-Exo70A1(WT)* or *35S::YFP-Exo70A1(S328A)*. For complementation studies, due to the developmental and reproductive defects observed in *exo70A1-1^−/−^* (28), we transformed heterozygous lines (*exo70A1-1^+/-^*), followed by screening for transformants that were homozygous for *exo70A1-1^−/−^* that harbored the transgene. These transgenic lines in the *exo70A1-1^−/−^* background were recovered at very low frequency. As expected, the wild-type *35S::YFP-Exo70A1(WT)* fully rescued the *exo70A1-1* pollination defects and other associated developmental phenotypes (Fig.3 C - E). In contrast, *35S::YFP-Exo70A1(S328A)* was incapable of rescuing any of the *exo70A1-1* phenotypes (Fig. 3 C - E). The *35S::YFP-Exo70A1 (S328A)/ exo70A1-1* lines were phenotypically similar to *exo70A1-1* plants, affected in pollen acceptance and produced shorter siliques (Fig. 3 C-E, Supplementary Fig. S9A). This strongly indicated that S328 is necessary for Exo70A1 function. To discern whether the presence of Serine 328 or its phosphorylation was necessary for full functionality of Exo70A1, we complemented *exo70A1-1* mutant with *35S::YFP-Exo70A1(S328D),* a phospho-mimic version of Exo70A1 (Fig. 3C-E, Supplementary Fig. S9B).The Exo70A1 phosophomimic was able to almost fully rescue all developmental and pollination phenotypes, further validating the requirement of S328 phosphorylation for Exo70A1 function (Fig. 3C-E, Supplementary Fig. S9B)

### Serine 328 of Exo70A1 is required for plasma membrane localization of Exo70A1

To examine the role of S328 phosphorylation on Exo70A1 localization, we observed localization of YFP tagged versions of Exo70A1 in *exo70A1-1* background (Fig. 4 A – D). YFP fluorescence from YFP-Exo70A1 (WT) could be readily visualized in the well-extended papillary cells of Arabidopsis stage 12 flowers. In unpollinated stigmas, YFP-Exo70A1(WT) localized to the plasma membrane as previously reported (18) (Fig. 4B). However, YFP-Exo70A1 (S328A) failed to localize to the plasma membrane and was found diffused throughout the cytoplasm (Fig. 4C). The lack of plasma membrane localization suggested that S328A mutation likely caused an aberration in trafficking and impaired its ability to localize at the plasma membrane where Exo70A1 controls exocytosis. This localization defect observed with the S328A form was fully rescued by the YFP-Exo70A1 (S328D) phosphomimic (Fig. 4D). The similarity in localization between Exo70A1(WT) and Exo70A1(S328D) provides strong evidence that phosphorylation of Exo70A1 at S328 is likely a necessary event for membrane localization of Exo70A1, where it can regulate exocytosis.

**Fig. 4.**
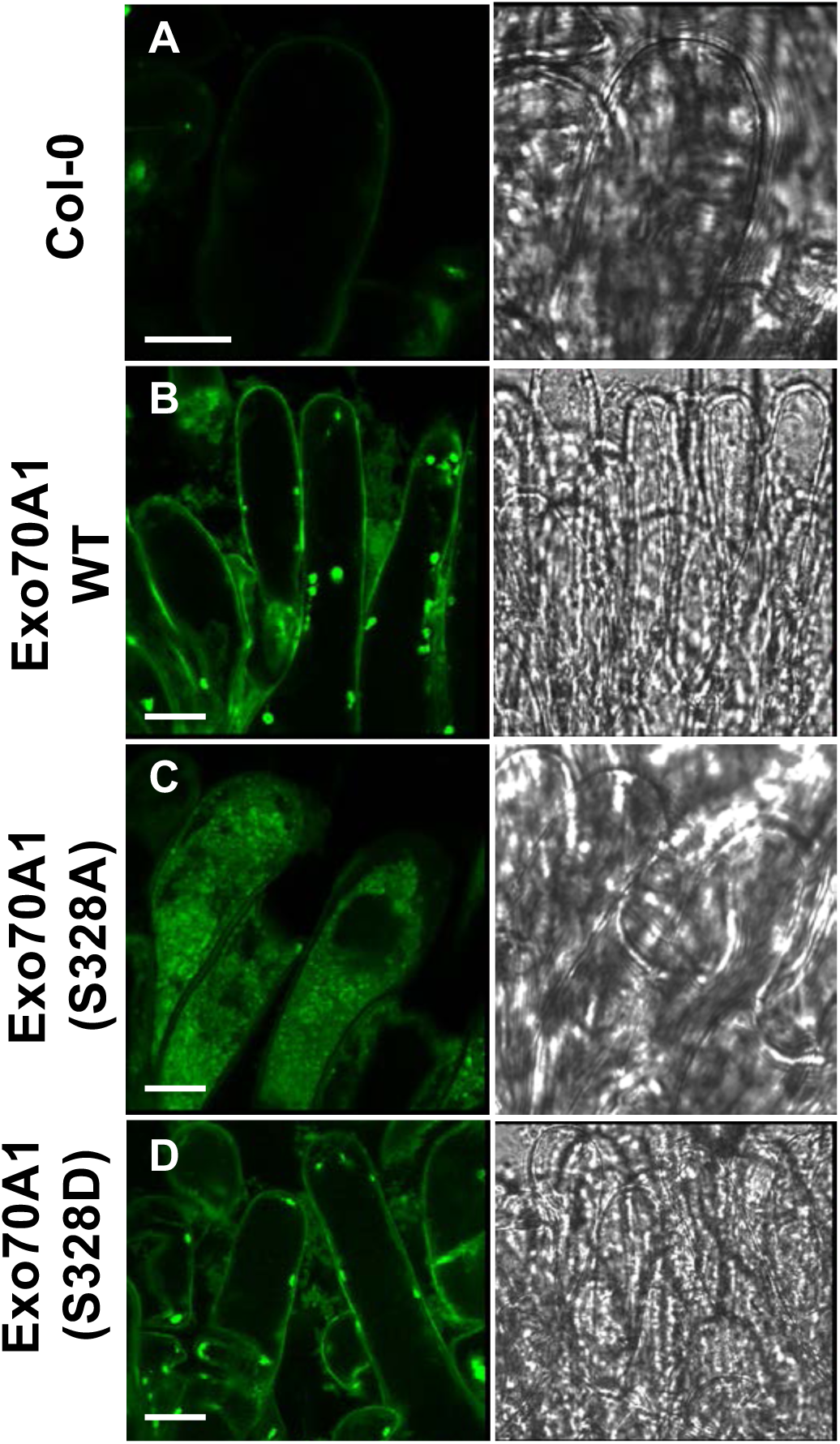
Disruption of Exo70A1 MAPK phosphorylation motif impairs its membrane localization. Confocal imaging of stage 12 papillary cells from Col-0 **(A)**, *35S::YFP-Exo70A1(WT)/exo70A1* **(B)** *, 35S::YFP-Exo70A1(S328A)/exo70A1 (Line# 2)* **(C)** and *35S::YFP-Exo70A1(S328D)/exo70A1 (Line# 41)* **(D)**. Membrane localization was changed to diffused and cytosolic in *35S::YFP-Exo70A1(S328A)/exo70A1* mutant whereas Phospho-mimic version 35S::YFP-Exo70A1(S328D)/exo70A1 restores the plasma membrane localization. Left column shows zoomed in images of papillary cells, right column shows bright field (BF) images of same papillary cells, scale: 5 µm.

## Discussion

Through this study, we were able to show that a subset of functionally redundant MAPK pathway components converges on the compatibility factor Exo70A1 to mediate stigma receptivity. Although the in-gel kinase assays detected three MAPKs, MPK3, MPK4 and MPK6 that were constitutively active in stage 12 stigmas of Arabidopsis, mutational analysis revealed that only MPK3 and MPK4 were required for pollination responses to occur. The loss of these two proteins resulted in severe reduction in pollen germination and pollen tube growth in spite of having homomorphic flowers that were unaffected in pollen transfer to the stigmas. Our assays had to rely on natural pollination as MAPKs are quite sensitive to manipulation and we were also utilizing the stigma-specific RNAi-mediated suppression of *MPK4* to circumvent lethality or severe phenotypes associated with the double knock-out mutants (23, 29, 30). Given that all these three kinases are commonly associated with defense against pathogen (7, 31–34), we could predict that one of the major roles of these kinases would be to protect the reproductive tissues from invading pathogens while simultaneously promoting stigma receptivity to compatible pollen. The constitutive accumulation of ROS in mature stigmas (7) suggests that it could be a candidate upstream signal as ROS are known to activate these kinases (35, 36).

We have identified that the MPK3/4 duo is regulated by five upstream MKKs. Loss of function or suppression of MKK1, MKK2, MKK3, MKK7 and MKK9 in the *mkk1/2/3Ri/mkk7/9* quintuple mutant led to reduced pollen attachment and the mutant phenocopied the *mpk4Ri/mpk3* pollination phenotype. The extreme functional redundancy observed at the MKK tier in facilitating stigma receptivity is certainly a paradigm shift in plant MAPK signaling where it is believed that fewer MKKs regulate all the MPKs (26, 37–40). Although most of the MKKs can interact with multiple downstream MPKs (25), this does not preclude the possibility of the *vice-versa* where a selected smaller subset of MPKs is regulated by multiple MKKs under certain conditions or in a specific tissue such as the stigmatic papillary cells. It is also plausible that since MPK3 and MPK4 belong to subgroups A and B respectively (41), an increased number of MKKs could interact with these kinases to influence stigma receptivity.

We have been able to show that the activated MPK3/4 pair converges on Exo70A1, a compatibility factor that regulates pollen hydration and germination in both Brassica and Arabidopsis (18, 42). Both MPK3 and MPK4 phosphorylate Exo70A1 at the consensus SP motif and this phosphorylation is essential to maintain stigma receptivity (Figure 3). A similar scenario has been observed in mammalian cells where ERK1/2-dependent phosphorylation of Exo70 was shown to facilitate interaction of Exo70 with other exocyst subunits to promote exocytosis in response to growth factor signaling (17). Both MPK3 and MPK4 also have TEY phosphorylation residues in their activation loop and belong to ERK1/2 class of mammalian kinases (43). Mutation in the consensus phosphorylation motif of Exo70A1 (S328A) disrupted its localization and was incapable of rescuing any of the *exo70A1-1* defects (Figure 3). Our phenotypic observations with the phosphodeficient Exo70A1 (S328A) and the phospohomimic Exo70A1 (S328D) clearly indicate the requirement of phosphorylation of Serine 328 for full functionality of Exo70A1 (Figure 3). Very similar to our observations, Exo70 (S250A) was unable to stimulate exocytosis in mammalian cells, while the phosphomimic was sufficient to rescue this defect (17). In Arabidopsis, mutating the MPK3/6-specific MAPK phosphorylation motif of the transcription factor, SPEECHLESS (SPCH) has been shown to cause over-proliferation of stomata (44). In metastatic melanoma cells, it was demonstrated that BRAF with V600E mutation causes constitutive activation of RAF-MEK-ERK pathways that converge on Exo70 phosphorylation leading to actin dynamics and secretion of matrix metalloproteases (45).

Although there are 23 paralogs in Arabidopsis, Exo70A1 is highly related to other eukaryotic Exo70s involved in exocytosis (28). Loss-of-function of Exo70A1 alone can cause a dwarf phenotype along with non-receptive stigmatic papillae (18, 28). Through an RNAi approach, each of the subunits of the complete octomeric complex was also shown to be required for conferring stigma receptivity, further validating the role of exocytosis as a crucial process during pollen acceptance (46). Exocyst subunits are known to dock at the plasma membrane (PM) and create sites for vesicle tethering and Exo70A1 is also required for plasma membrane localization of other subunits such as Sec6 (47). The lack of PM localization of Exo70A1 in the absence of S328 phosphorylation suggests that phosphorylation could result in interaction of Exo70A1 with other subunits to traffic to the plasma membrane where it can regulate exocytosis. Alternatively, phosphorylation of Exo70A1 could recruit it to the membrane where it allows the assembly of exocyst complex through associating with other subunits to facilitate vesicle tethering and exocytosis. Consistent with this, Loss of function of Exo70A1 in Arabidopsis and *Brassica napus* resulted in accumulation of these vesicles in the cytoplasm of both unpollinated and pollinated papillary cells (42). Accumulation of these vesicles in the cytoplasm is proposed to restrict the necessary resources from being delivered to the germinating pollen (42).

Collectively, we have been able to unravel a novel role for MAPK signaling during pollination that converges on a key regulator of the stigmatic exocytotic pathway essential for stigma receptivity (Figure 5). The extreme functional redundancy observed at the MKK level of the cascade breaks the current paradigm where a smaller number of MKKs are believed to regulate a large class of MPKs, suggesting that size of gene family may not necessarily correlate with overlapping functions. Given the agronomic importance of various crop plants that belong to Brassicaceae, understanding the signaling mechanism during compatible pollination responses could lead to developing strategies to improve crop yield.

**Fig. 5.**
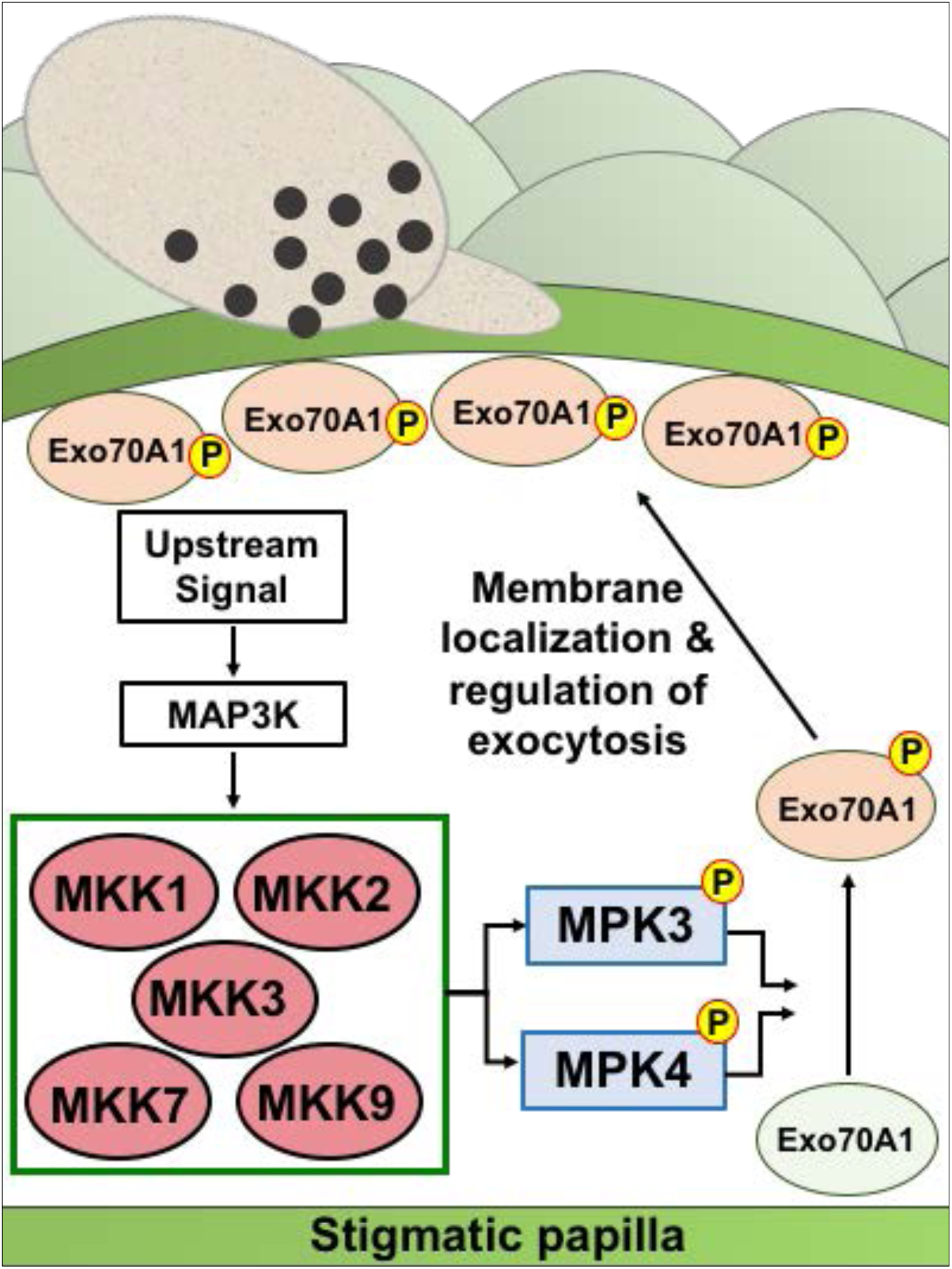
Proposed model for signaling during compatible pollination. In stigmatic papillary cells, MAPK cascade phosphorylates Exo70A1 to localize at plasma membranes where it regulates exocytosis to facilitate successful pollination.

## Materials and Methods

### Plant materials and growth conditions

*Arabidopsis thaliana* Col-0 ecotype was used as wild-type. Mutant alleles of *mpk3* (SALK_151594), *mpk6* (SALK_073907), *mkk7* (SM_CS110477), *mkk9* (SAIL_60_H_06) and *exo70A1-1* (SALK_014826) were obtained from ABRC and homozygous lines were isolated. One of the single mutant *mpk3-DG* (22)was obtained from Dr. Brian Ellis (University of British Columbia, Vancouver, Canada). All seeds were surface sterilized and plated on 0.5 X MS, 1% sucrose and 0.7% phytagar, stratified for 3 - 5 days at 4 °C in dark and germinated in plant growth incubators. The seedlings were then transplanted to soil and plants were maintained in growth chambers at 22 °C with a 16 h / 8 h (light / dark) cycle and a light intensity of 120 μmol m^−2^ s^−1^ 7 days.

### RNAi constructs and generation of multi-locus mutants

RNAi constructs of *MPK3, MPK4* and *MPK6* were generated using the first 600 bases of the ORF as previously described (18). Infusion cloning (Clontech) was used to introduce these RNAi constructs into modified pBin19 binary vector containing stigma-specific *SLR1* promoter, to drive suppression of these genes exclusively in the stigmatic tissue. The primers used for developing RNAi constructs are shown in supplemental Table 3. For generating RNAi constructs of *MKKs*, tandem constructs of *MKK1/2*, *MKK1/2/3* and *MKK4/5,* were synthesized (GenScript, Piscataway, NJ, USA), using the first 200 - 300 bases of the ORF of each gene. These constructs were then utilized for building tandem RNAi constructs of *MKK1/2*, *MKK1/2/3* and *MKK4/5* through infusion cloning in pBin19 vector (for primers refer to supplemental Table 3). *AtExo70A1* ORF was amplified and cloned in pGEX4T-1. Site directed mutagenesis was used to change serine 328 to alanine (S328A) and aspartic acid (S328D) by quick change PCR (Invitrogen). Clones were sequenced to confirm the changes and absence of mismatches. *Exo70A1*, *Exo70A1 (S328A)* and *Exo70A1 (S328D)* were PCR amplified and cloned as a C-terminal fusion with *YFP* in pCAMBIA 1301 vector to generate *35S::YFP-Exo70A1*, *35S::YFP-Exo70A1 (S328A)* and *35S::YFP-Exo70A1 (S328D)* constructs. Primers used for generating these constructs are shown in supplemental Table 4. Col-0 and other mutant plants were transformed using *Agrobacterium tumefaciens* (GV3101)-mediated transformation through floral dip method (48). Transformant selection was performed by screening on 0.5 X MS, 1 % sucrose plates containing either hygromycin (25 mg/L) or kanamycin (50 mg/L). Putative transformants were then transplanted to soil, maintained in growth chambers and subjected to further analysis.

### Recombinant protein production and purification

GST fusion clones of *CAKK1* and *CAKK7* were obtained from Dr. Brian Ellis (University of British Columbia, Vancouver, Canada). MPK3 and MPK4 ORFs were cloned under 6xHIS tag in pET15b vector. All recombinant glutathione S-transferase (GST) fusion and 6xHIS proteins were expressed in *Escherichia coli* BL21 cells. Cultures were induced with either .01 or 0.1 mM isopropylthio-galactoside (IPTG) for 6 h at 30 °C, followed by purification according to the manufacturer’s protocol (Sigma-Aldrich).

### In-gel kinase assay

In-gel kinase assays were performed as described earlier (9). Stage 12 stigmas from the various genotypes were collected in liquid nitrogen and stored at – 80 °C until further analysis. For protein extraction, stigmas were ground in extraction buffer (100 mM Hepes, pH 7.5, 5 mM EDTA, 5 mM EGTA, 10 mM DTT, 10 mM Na_3_VO_4_, 10 mM NaF, 50 mM β - glycerophosphate, 1 mM phenylmethylsulfonyl fluoride (PMSF) and protease inhibitor cocktail (Roche)), centrifuged at 13000 rpm for 15 min at 4 °C to pellet the debris. Extracts containing 5 μg of proteins were resolved on a 10% SDS gel embedded with 250 μg/mL of myelin basic protein (MBP). SDS was removed by washing the gel with washing buffer (25 mM Tris, pH 7.5, 0.5 mM DTT, 0.1 mM Na_3_VO_4_, 5 mM NaF, 0.5 mg/mL BSA, 0.1 % Triton X-100 (V/V)) for three times at room temperature. The gel was then incubated in renaturation buffer (25 mM Tris, pH 7.5, 1 mM DTT, 0.1 mM Na_3_VO_4_ and 5mM NaF) overnight at 4 °C with three buffer changes. The gel was then transferred to reaction buffer (25 mM Tris, pH 7.5, 2 mM EGTA, 12 mM MgCl_2_, 1 mM DTT, 0.1 mM Na_3_VO_4_, 200 nM ATP) for 30 min at room temperature followed by 60 min incubation in same reaction buffer with 25 μCi γ ^32^P (specific activity 3000 Ci/mmol). After incubation gel was transferred to 5% trichloroacetic acid (TCA) (w/v) and 1% NaPPi (w/v) to stop the reaction. The gel was washed for 5 times in one hour to remove unincorporated γ ^32^P followed by drying in Gel Air dryer for 90 min with heating and 30 min at room temperature. The dried gel was subjected to phosphor-imaging screen to detect kinase activity.

### *In vitro* kinase assay

Recombinant purified proteins were used for *in vitro* phosphorylation assay. Recombinant 6xHIS-MPK4 (5μg) was activated with GST-CAKK1 (1μg) in phosphorylation reaction buffer (20 mM Hepes, pH 7.5, 10mM MgCl_2_, 1 mM DTT) at 28 °C for 1.5 h. Activated proteins were then used to phosphorylate recombinant GST-Exo70A1 (5 and 10μg) in same reaction buffer with the addition of 25 μM ATP and γ ^32^P-ATP (1 μCi per reaction specific activity 3000 Ci/mmol) for 30 min at 28 °C. The reaction was stopped by adding 6X SDS loading buffer. Reaction mixture was resolved on 10% SDS gel followed by staining and destaining of gels to visualize proteins. The gel was air dried for 90 min with heating and 30 min at room temperature. Phosphorylated GST-Exo70A1 was visualized by autoradiography.

### Dot Blot Analysis

AtExo70A1 short peptides from residue 323 to 335 (AR***<u>S</u>***KR***<u>S</u>***PEKLFVL and AR***<u>S</u>***KR***<u>A</u>***PEKLFVL) were synthesized from GL Biochem, Shanghai, China. Peptides were dissolved in 1X PBS (pH 7.0 to 7.5) and immobilized on PVDF membrane in various concentrations followed by air drying for 2 h. Membranes were blocked with 5% BSA (w/v) for 1 hr and incubated in kinase assay buffer (20 mM Hepes, pH 7.5, 10mM MgCl_2_) at 30°C with either CAKK1-6xHIS-MPK4 or CAKK7-6XHIS-MPK3 in the presence of phosphatase inhibitor (25 mM NaF, 0.2 M NaPPi and 2 mM Na3VO4). Membranes were probed with anti pSer antibody 1:5000 (ab9332 Abcam) in the presence of phosphatase inhibitors (25 mM NaF and 1 mM Na3VO4) followed by goat anti-rabbit secondary antibody and development with ECL (Amersham).

### Aniline blue assay

Aniline blue assay was performed as described earlier (18). In case of inhibitor analysis, Col-0 stigmas were incubated in U0126 overnight and pollinated for 4 h with Col-0 pollen grains. Arabidopsis Col-0 and mutants were allowed to naturally pollinate and stigmas were collected 4 h after flower opening, fixed in 3:1 ethanol: glacial acetic acid for 30 min followed by 1 h incubation in 1 N NaOH at 60 °C. Stigmas were then washed with distilled water for 3 times and stained for 30 min with basic aniline blue (0.1% aniline blue in 0.1 M K3PO4). Stained stigmas were mounted in 50% glycerol and pollen attachment and pollen tube germination were observed in the blue channel under Leica DMR epifluorescence microscope.

### Reverse transcriptase quantitative PCR

Stage 12 stigmas from Col-0 and the various mutants were collected in liquid nitrogen and stored at – 80 °C until further analysis. Total RNA was extracted from these stigmas using RNeasy columns with plant RNA extraction aid (Qiagen RNeasy Mini Kit) according to manufacturer instructions. RNA was quantified with NanoDrop spectrophotometer and RNA integrity was further confirmed by resolving on an agarose gel. Total RNA was treated with DNase I (Thermo Scientific Bio) and reverse transcription was performed with 500 ng of RNA using superscript II (Invitrogen) according to manufacturer’s instructions. Quantitative PCRs (qPCR) were performed with SybrGreen dye in StepOnePlus PCR detector (Applied Biosystems). PCR amplification was performed for 40 cycles at 95 °C for 3 s and 60 °C for 30 s with a preceding initial enzyme activation of 20 s at 95 °C. Relative expression levels were calculated by delta-delta-Ct method. *UBQ10* was used as endogenous control for all samples. Primers used for qPCR for *MPK3, MPK4, MKK1, MKK2, MKK3, MKK7* and *MKK9* are shown in supplemental Table 5.

### Confocal microscopy

Leica SP5 laser confocal microscope was used for Exo70A1 localization studies. Images were acquired with 40 X oil-immersion objective and the highly sensitive Leica HyD detector. Arabidopsis 35S::YFP-Exo70A1 and 35S::YFP-Exo70A1 (S328A) and 35S::YFP-Exo70A1 (S328D) stage 12 stigmas were mounted in 50% glycerol and subjected to confocal imaging. To capture the YFP signal, stigmas were scanned with argon laser (Excitation 476 nm: emission 490-535nm). Autofluorescence was detected between 590-720nm emission wavelength and merged images were generated by combining both channels. At least three different transgenic lines were examined for each construct.

## ACKNOWLEDGMENTS

We thank Dr. Brian Ellis (UBC) for sharing recombinant MPK and MKK constructs. We thank Dr. Douglas Muench for the epi-fluorescence microscope facility. This work was supported by Natural Sciences and Engineering Research Council of Canada funding for MAS.

## Author contributions

M.J., and M.A.S initiated the project. M.J., S.S., and M.A.S. designed research; M.J., S.S., K.A. and M.A.S. performed all the experiments; M.J. and M.A.S. analyzed data and wrote the manuscript. All authors were involved in manuscript discussions and commented on the manuscript.

## Supplemental Information

### Supplemental Figure Legends

**Fig S1.**
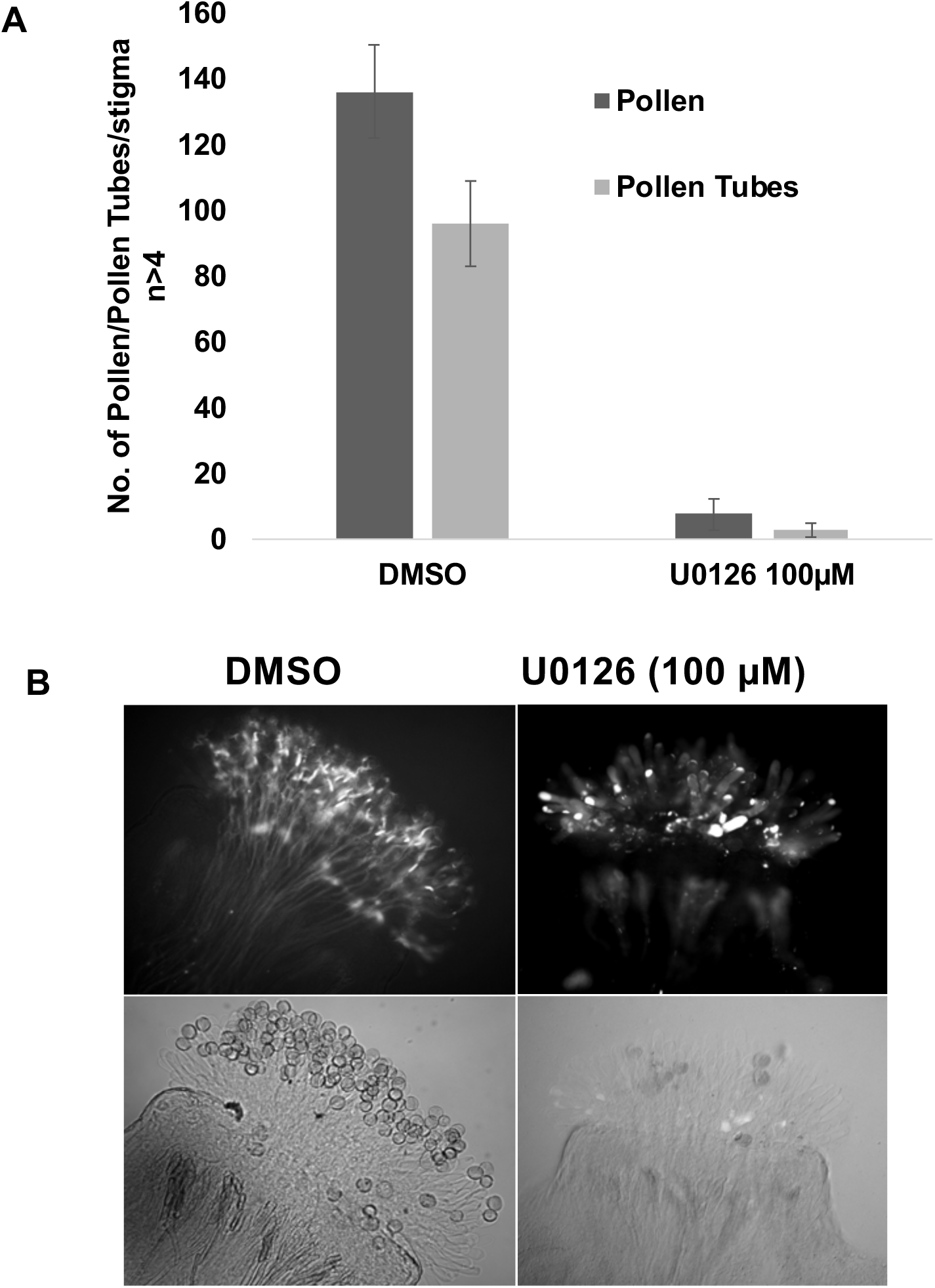
Inhibition of MAPKK(s) in Arabidopsis stigmas compromises pollination. *Arabidopsis thaliana* flowers were used for this assay. Arabidopsis stigmas were incubated overnight in 100 μM U0126, pollinated for 4 h with Col-0 pollen and stained with aniline blue to assess pollination. **(A)** Graph representing average number of pollen attached and pollen tubes germinated following either DMSO treatment or U0126 treatment (n>7). Error bars indicate ± SE. Asterisk represents values significantly different from DMSO treatment at P<0.05. **(B)** Aniline blue image of stigma showing reduced pollen attachment and pollen tube germination following U0126 treatment.

**Fig S2.**
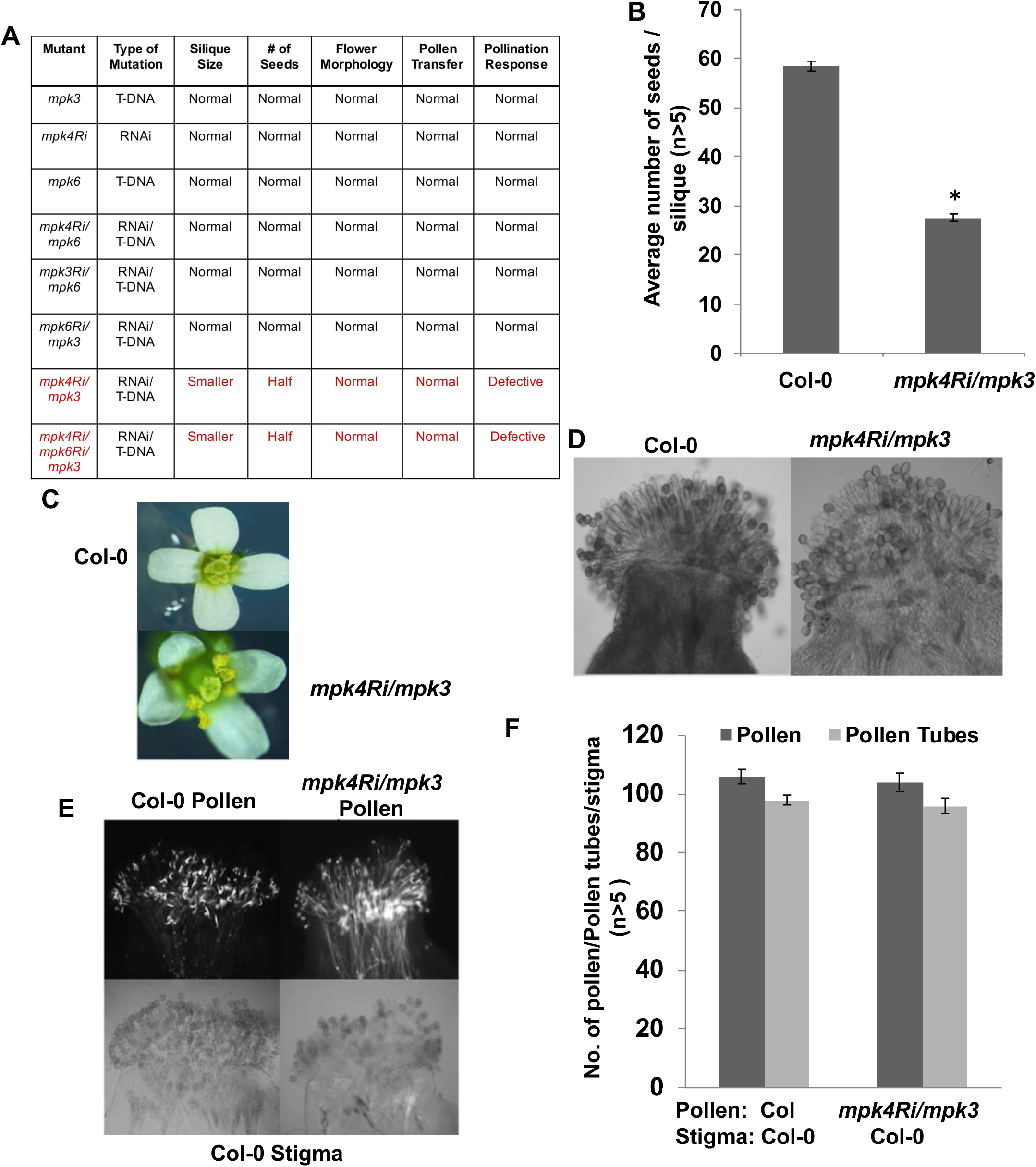
Pollination phenotypes of different individual and combinatorial *MAPK* class mutants. Floral development, pollen transfer and viability of *mpk4Ri/mpk3* mutant. **(A)** Table represents different mutants tested for various parameters and pollination phenotypes. **(B)** Graph representing the average number of seeds/silique in Col-0 and *mpk4Ri/mpk3* mutant (n>5). Error bars indicate ± SE. Asterisk indicate significant differences (*P<0.05). **(C)** Flowers of *mpk4Ri/mpk3* mutant are homomorphic and show no developmental defect. **(D)** Col-0 and *mpk4Ri/mpk3* mutant stigmas showing pollen grains naturally transferred on to papillary cells. Stigmas were collected from stage 13 open flowers 4 h after flower opening. **(E)** Reciprocal cross showing normal pollen viability of *mpk4Ri/mpk3* pollen grains. Col-0 stigma were hand pollinated with Col-0 and *mpk4Ri/mpk3* pollen grains for 4 h and subjected to aniline blue staining. **(F)** Graph represents average number of pollen attached and pollen tubes germinated per stigma following 4 h of pollination (n>5). Error bars indicate ±SE.

**Fig S3.**
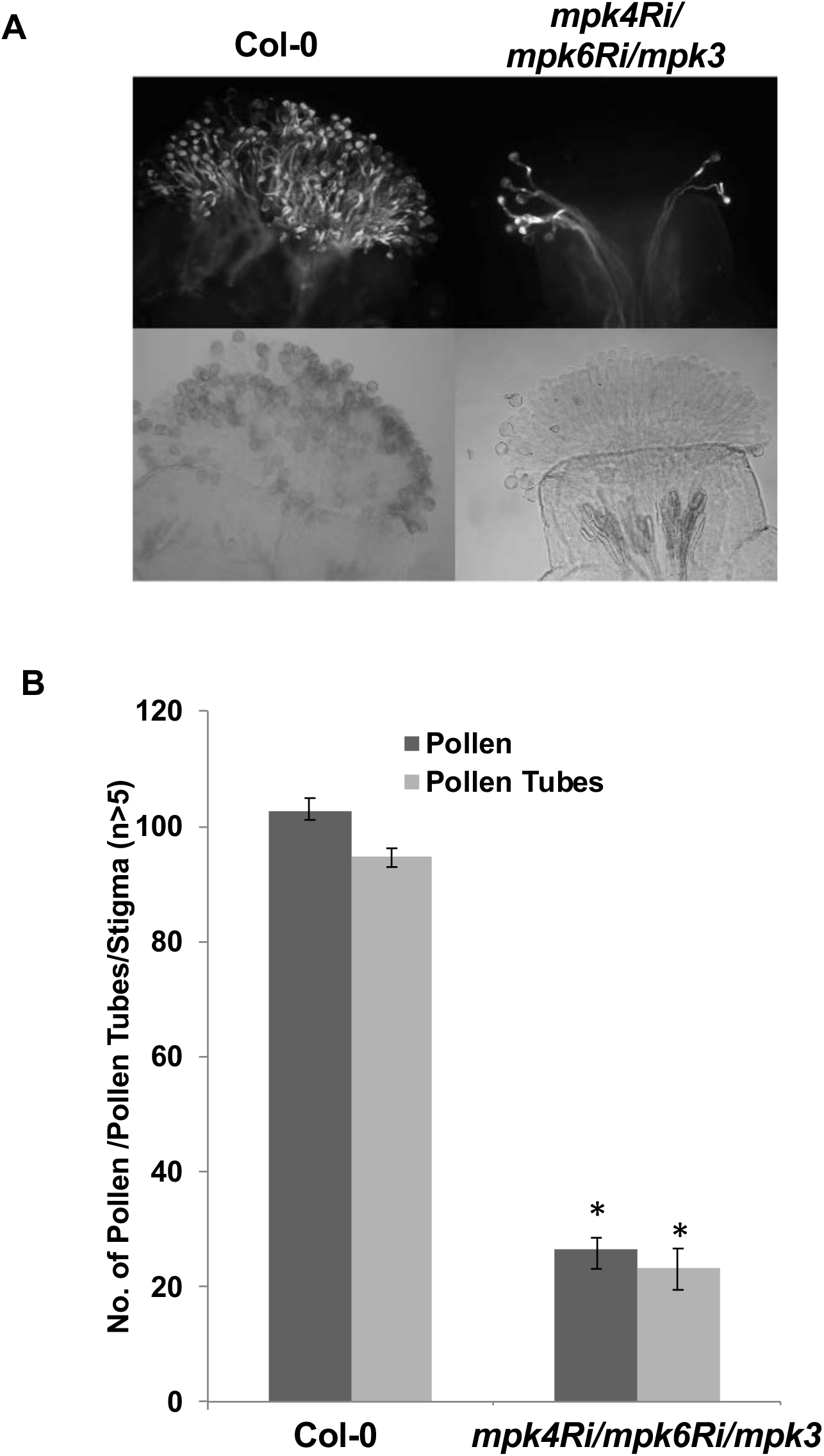
Genetic evidence for lack of a role for *MPK6* during pollination. Pollination defect in *mpk4Ri/mpk6Ri/mpk3* was similar to the double mutant *mpk4RNAi/mpk3*. **(A)** Aniline blue image showing pollen grains adhered and pollen tubes germinated on triple mutant stigma. Stigmas were pollinated for 4 h and subjected to aniline blue staining. **(B)** Graph represents the average number of pollen attached and pollen tubes germinated following 4 h of pollination. Asterisk represents values significantly different from Col-0 at P<0.05 (n>5). Error bars indicate ± SE.

**Fig.S4.**
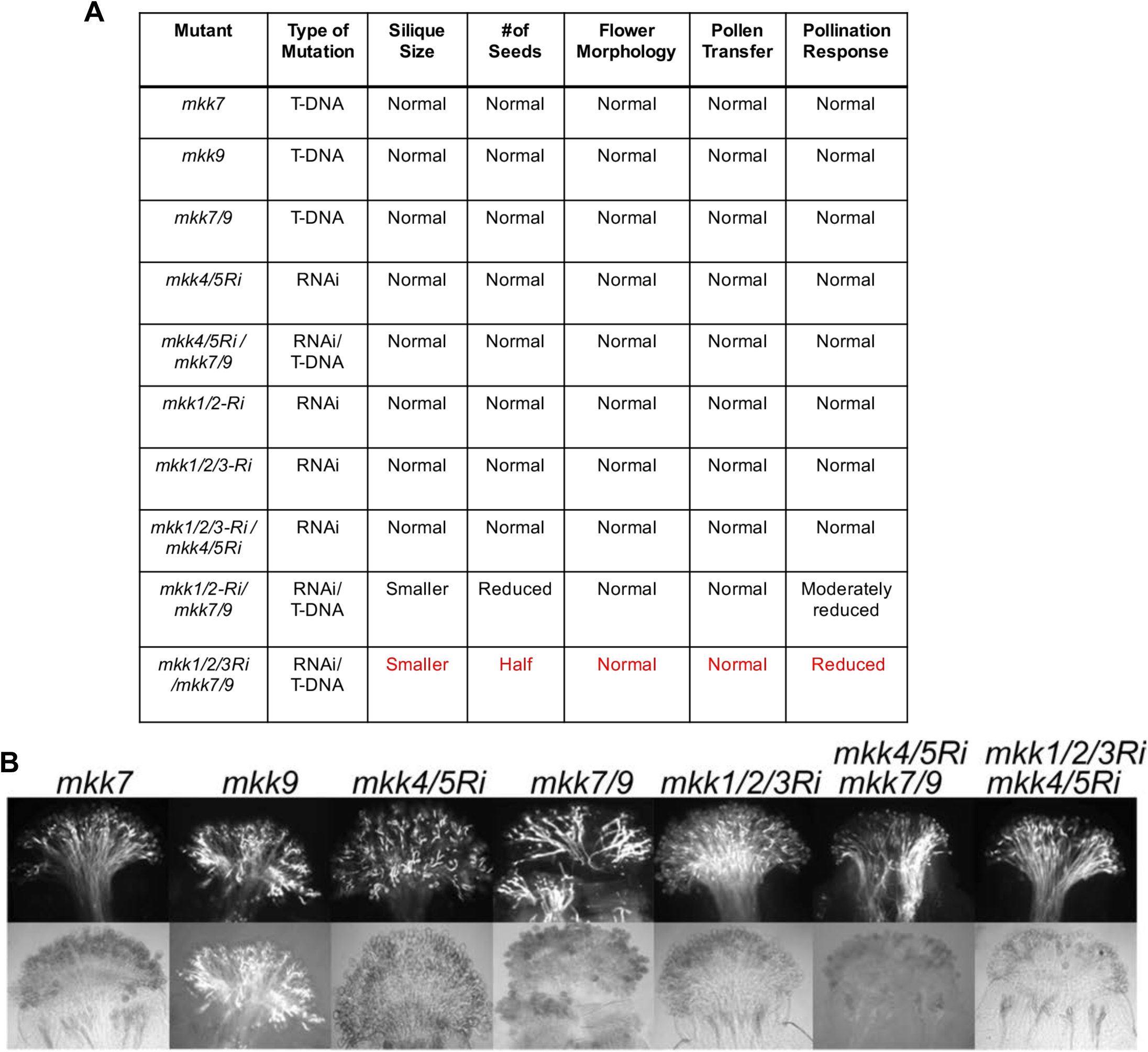
Five MKKs regulate compatible pollination. **(A)** Table represents different mutants tested for various parameters and pollination phenotypes. Quadruple mutant *mkk1/2Ri/mkk7/9* displayed partial breakdown of pollination responses while quintuple mutant *mkk1/2/3Ri/mkk7/9* phenocopied *mpk4Ri/mpk3* pollination defect. **(B)** Aniline blue assays of stigmas from various *MKK* single and combinatorial mutants showing pollen attachment and pollen tube growth following self-pollination. All stigmas were allowed to naturally pollinate and collected 4 h after flower opening. All RNAi constructs were placed under stigma-specific *SLR1* promoter.

**Fig S5.**
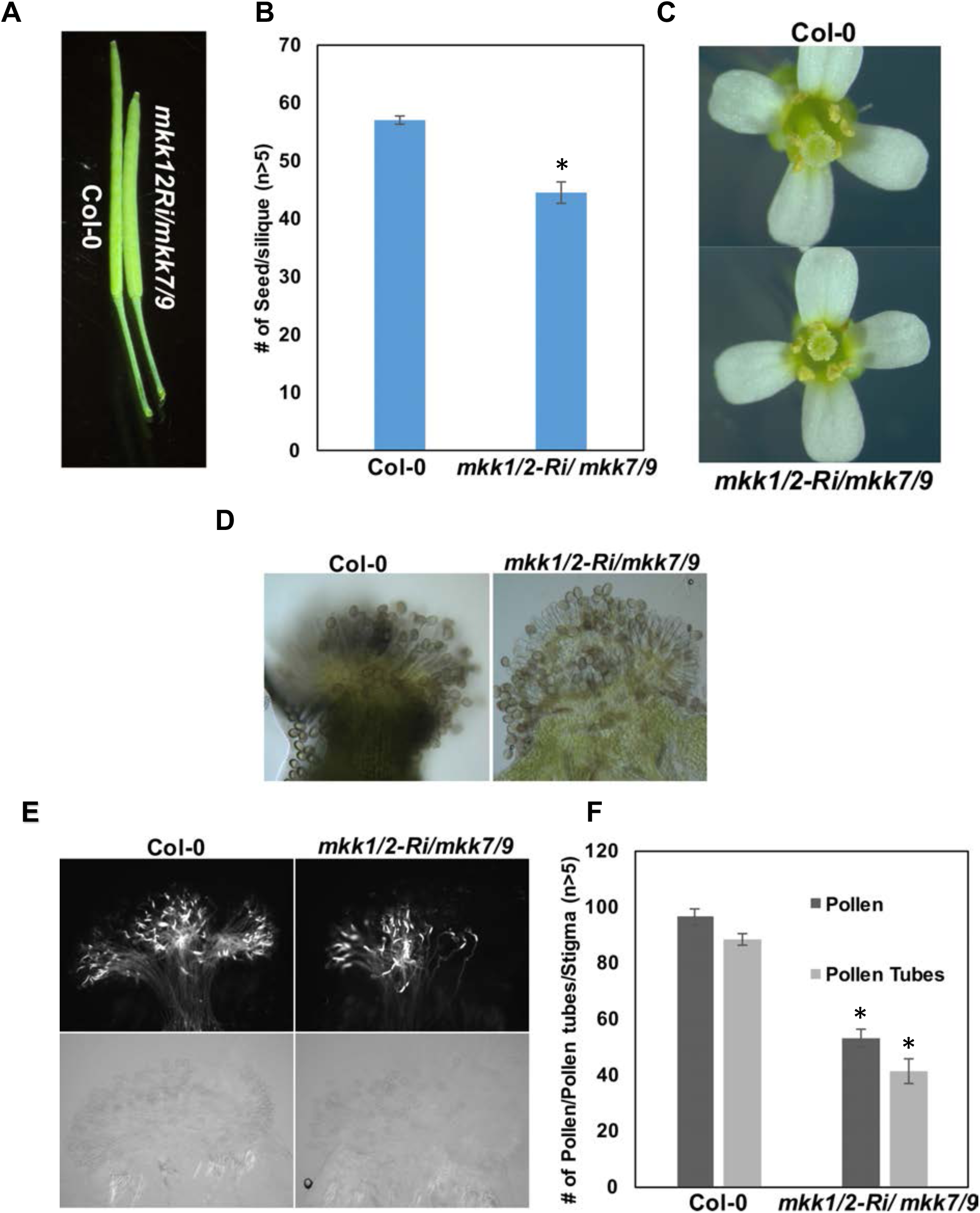
Suppression of *MKK1/2/7/9* results in partial defect in pollination. **(A)** Difference in silique sizes between Col-0 and *mkk1/2Ri/mkk7/9.* **(B)** Graph represents the average number of seeds in each silique. Error bars represent standard error of mean (n>5). Asterisks indicate significant differences (*P<0.05). **(C, D)** Mutant *mkk1/2Ri/mkk7/9* lines produce flowers of wild-type appearance and show no change in pollen transfer. **(E)** Aniline blue assays represent that moderate reduction in pollen attachment and germination observed in *mkk1/2Ri/mkk7/9* mutant lines compared to Col-0. **(F)** Graph represents average number of pollen attached and pollen tubes germinated per stigma following 4 h of pollination. Error bars indicate ± SE (n>5). Asterisks indicate significant differences (*P<0.05).

**Fig S6.**
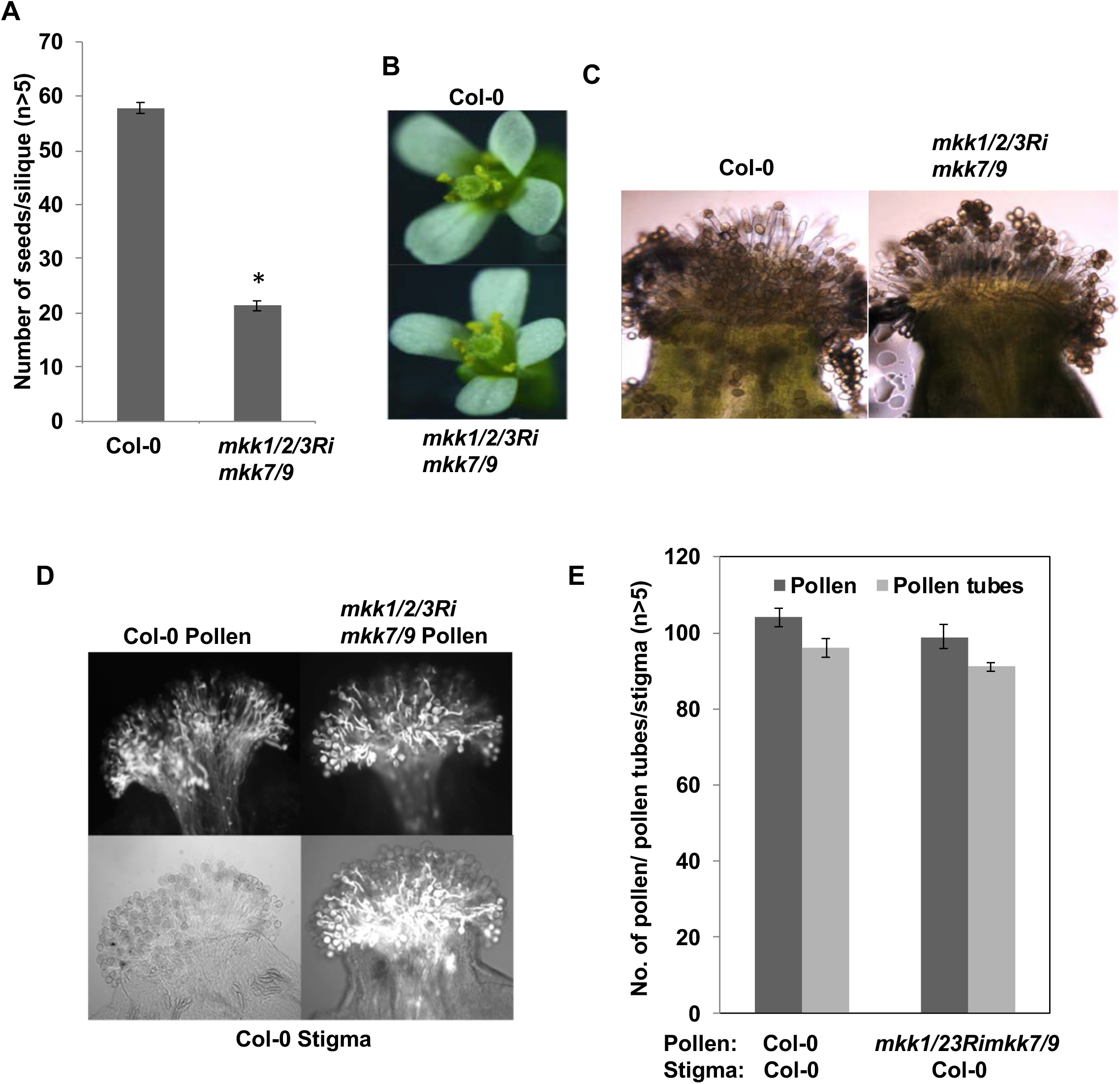
Suppression of five MKKs, *MKK1/2/3/7/9* results in pollination defects similar to *mpk4Ri/mpk3* mutant but does not alter floral architecture or pollen viability. **A)** Graph represents the average number of seeds/silique in Col-0 and quintuple *(mkk1/2/3Ri/mkk7/9)* mutant. Error bars indicate ±SE (n>5). Asterisk indicate significant differences (*P<0.05). **(B)** Col-0 and *mkk1/2/3Rimkk7/9* flowers after anthesis, mutant flowers are indistinguishable from Col-0. **(C)** Pollen transfer on stigma in WT Col-0 and *mkk1/2/3Ri/mkk7/9* mutant. No defect in pollen transfer efficiency was observed in mutant lines. **(D)** Reciprocal cross showing normal pollen viability of *mkk1/2/3Ri/mkk7/9* pollen. Col-0 stigmas were hand pollinated with Col-0 and *mkk1/2/3Ri/mkk7/9* pollen grains for 4 h and subjected to aniline blue staining. **(E)** Graph represents average number of pollen adhered and pollen tubes germinated per stigma (n>5). Error bars indicate ±SE.

**Fig S7.**
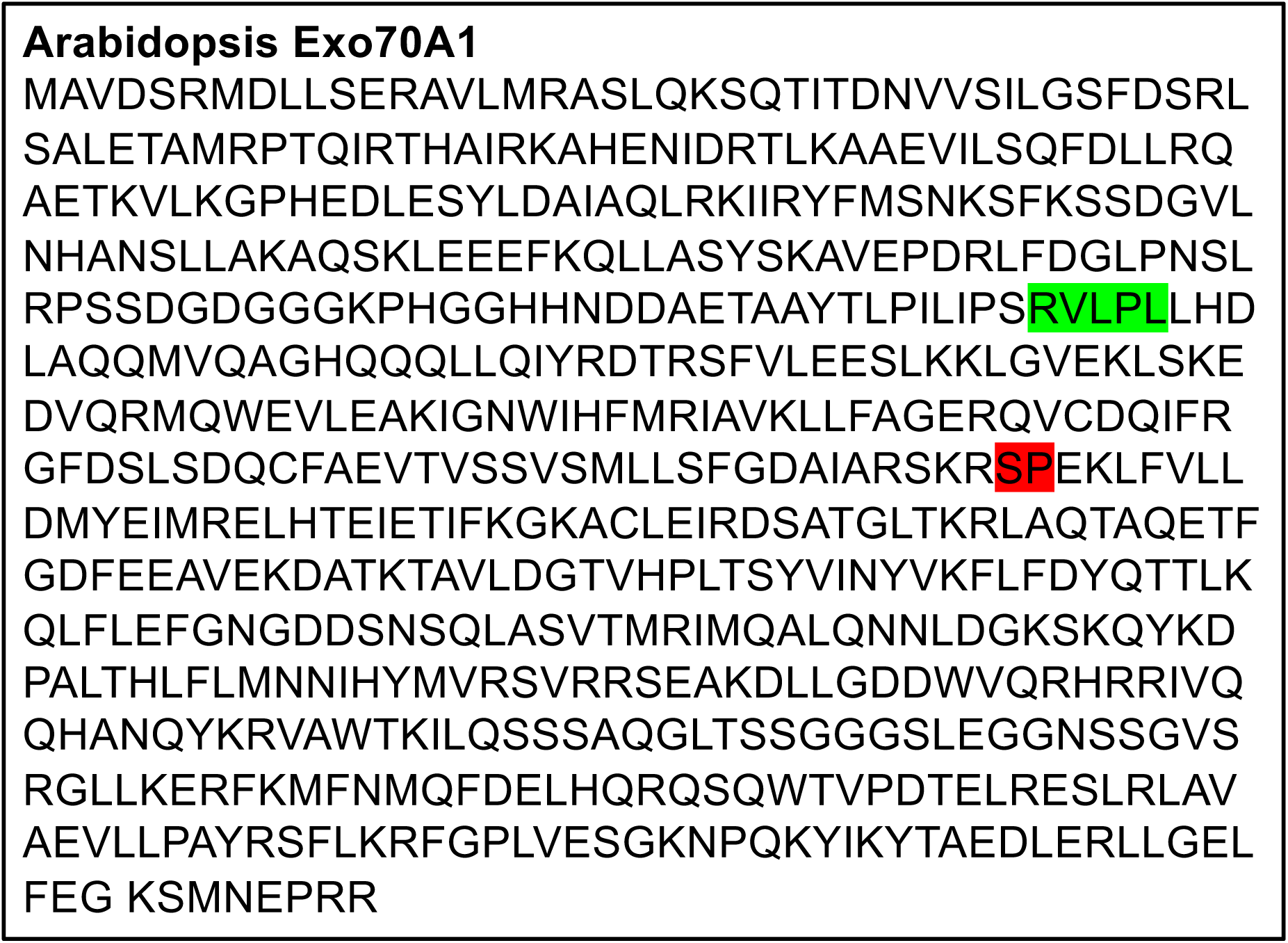
Protein Sequence of AtExo70A1. MAPK(s) phosphorylation consensus motif (S/TP) in Exo70A1 is highlighted in red. Docking domain for MAPK(s) is shown in green.

**Fig S8.**
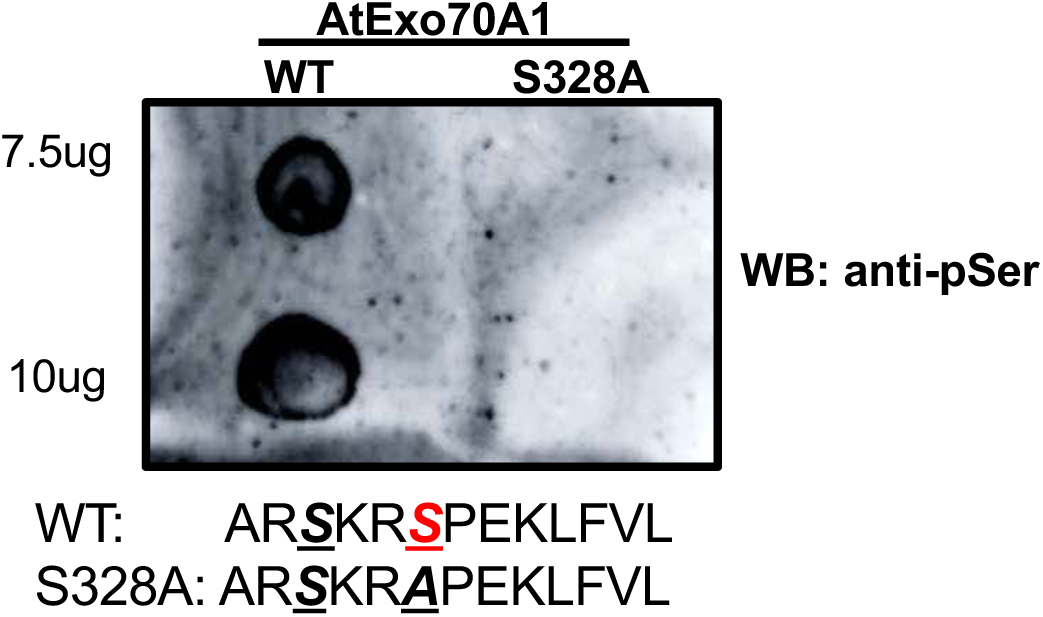
EXO70A1 is phosphorylated by MPK4 *in-vitro*. *In vitro* kinase assay showing phosphorylation after dot blotting of WT Exo70A1 peptide (residues 323-335) and S328A Exo70A1 peptide (residues 323-335). Kinase reaction was performed in the presence of GST-CAKK7 and 6xHIS-MPK3 and probed with anti-pSer antibody.

**Fig. S9.**
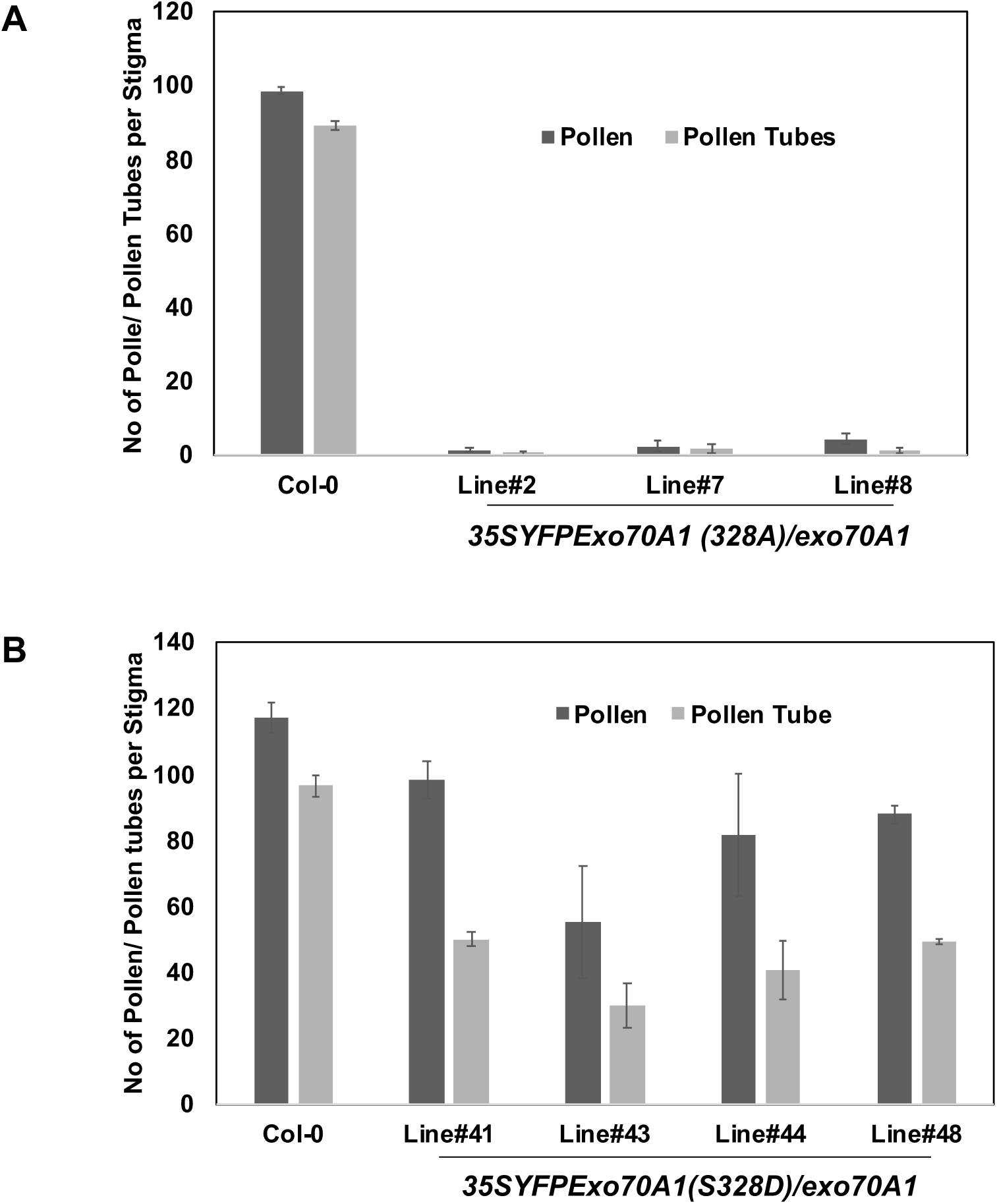
Serine 328 of Exo70A1 is required for full functionality of Exo70A1 during pollination. **(A)** Various transgenic lines harboring *35S::YFP-Exo70A1(S328A)* in the *exo70A1-1* background were subjected to pollination assays using Col-0 pollen. Graph represents the number of pollen attached and pollen tubes germinated. **(B)** Average number of Col-0 pollen attached and pollen tubes germinated on stigmas of different transgenic lines expressing *35S::YFP-Exo70A1(S328D),* the phosphomimic version of Exo70A1 in *exo70A1-1* background.

**Supplemental Table 1:**
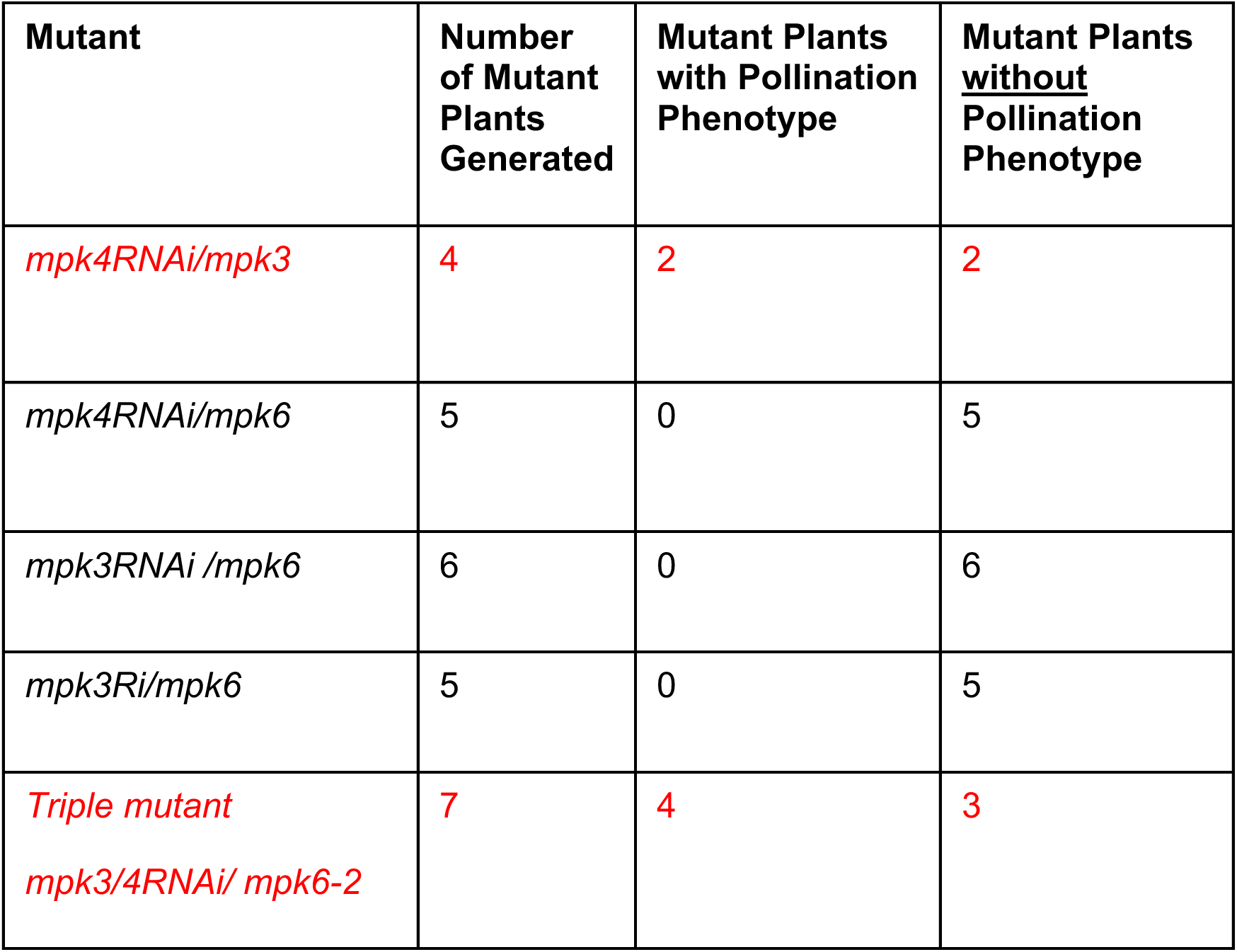
List of various double and triple MAPK mutants generated. Table represents the total number of mutant lines generated for each combination and number of mutant lines exhibiting pollination defects.

**Supplemental Table 2:** List of various combinatorial MAPKK mutants generated. Table shows total number of mutants created for various MKK combinations and their respective pollination defects.

| **Mutant** | **Number of Mutant Plants Generated** | **Mutant Plants with Pollination Phenotype** | **Mutant Plants <u>without</u> Pollination Phenotype** |
| --- | --- | --- | --- |
| *mkk4/5RNAi* | 9 | 0 | 9 |
| *mkk1/2/3RNAi* | 7 | 0 | 7 |
| *Quadruple*<br>*mkk1/2RNAi/mkk7/9* | 5 | 3 (Partial phenotype) | 2 |
| *mkk4/5RNAi/mkk7/9* | 7 | 0 | 7 |
| *Quintuple*<br>*mkk1/2/3/RNAi/mkk7/9* | 5 | 2 | 3 |
| *mkk1/2/3RNAi/mkk4/5RNAi* | 6 | 0 | 6 |

**Supplemental Table 3:**
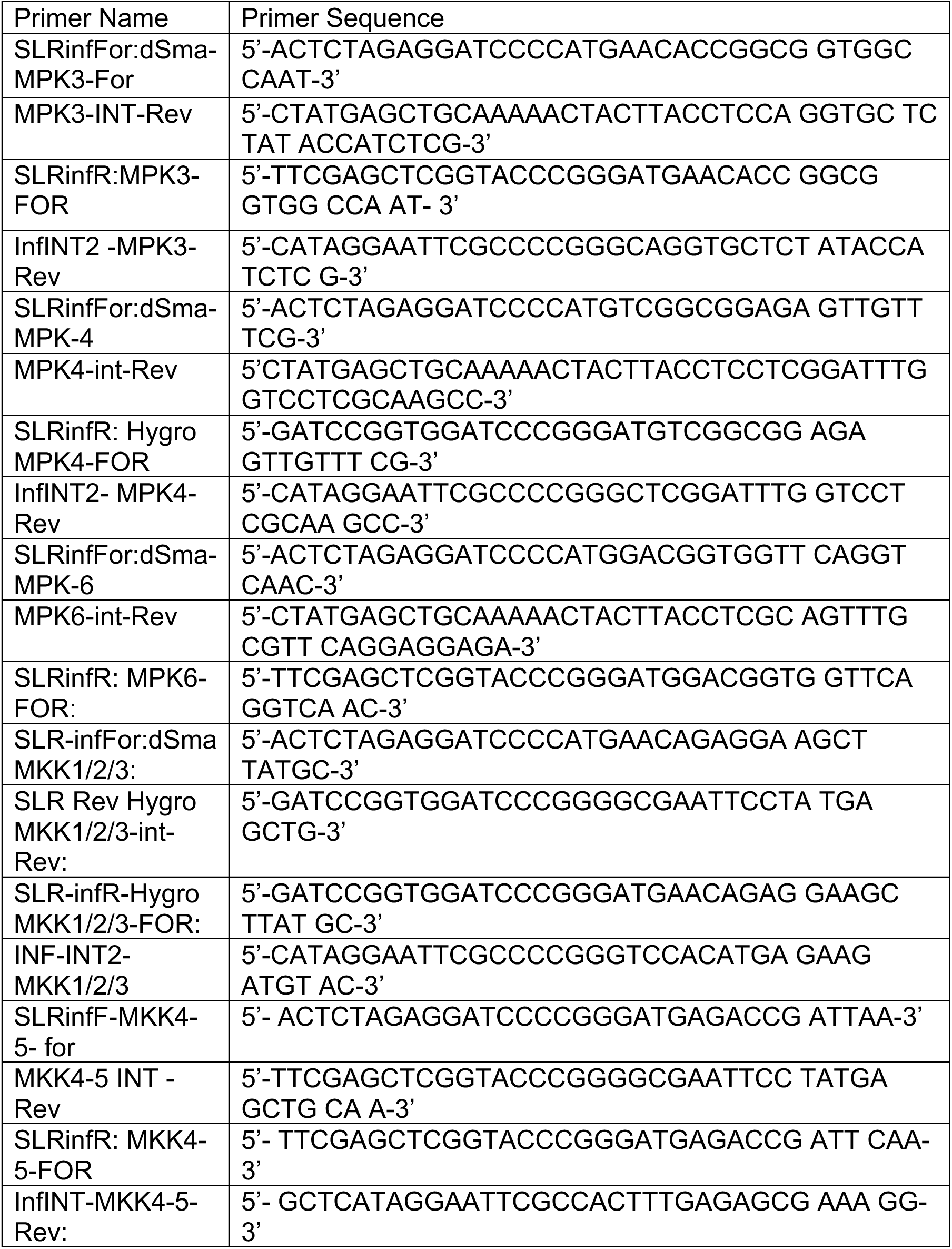
Primer sequences used for RNAi cloning construct.

**Supplemental Table 4:**
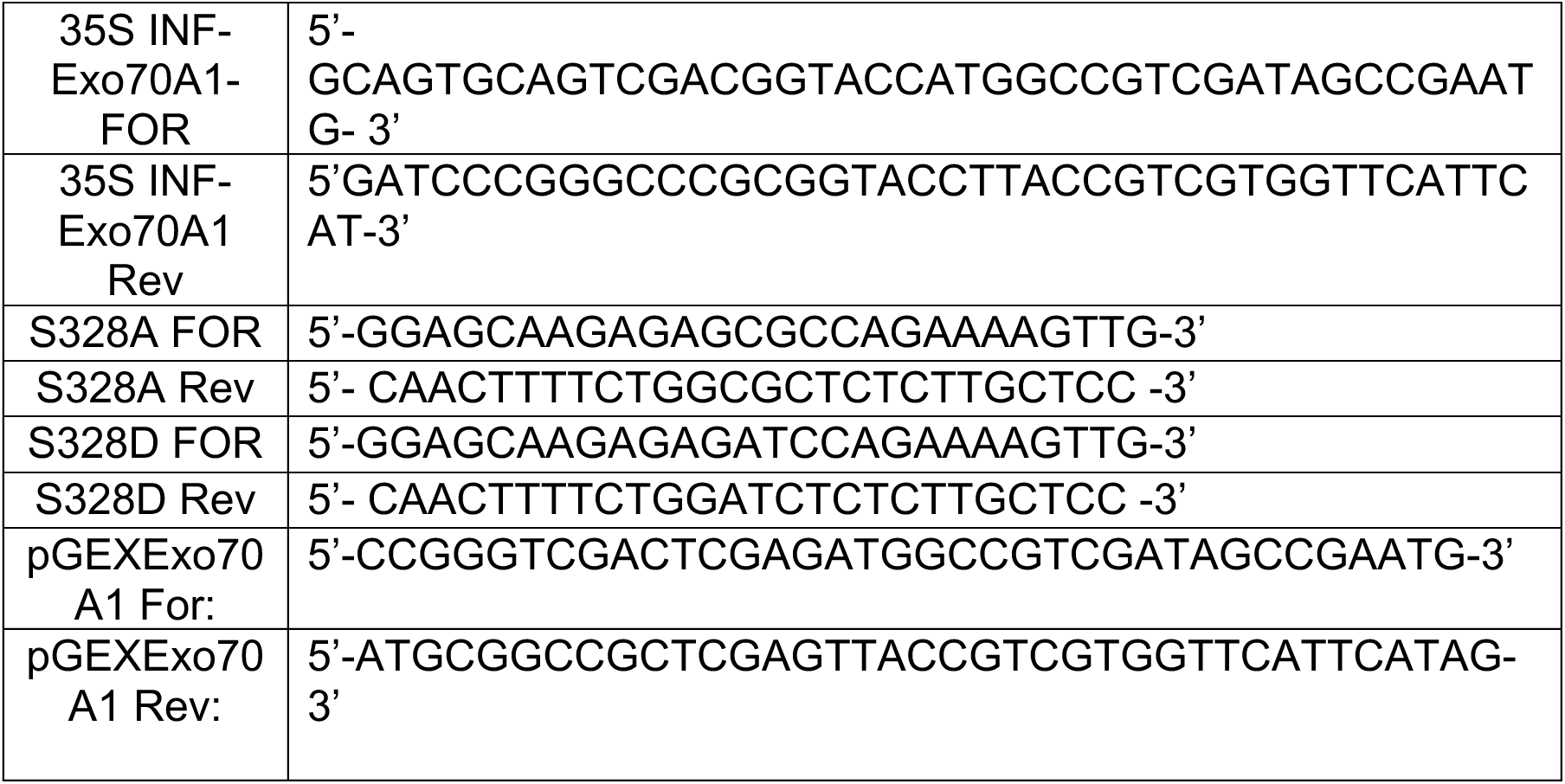
Primers used for *AtEx070A1* cloning.

**Supplemental Table 5:**
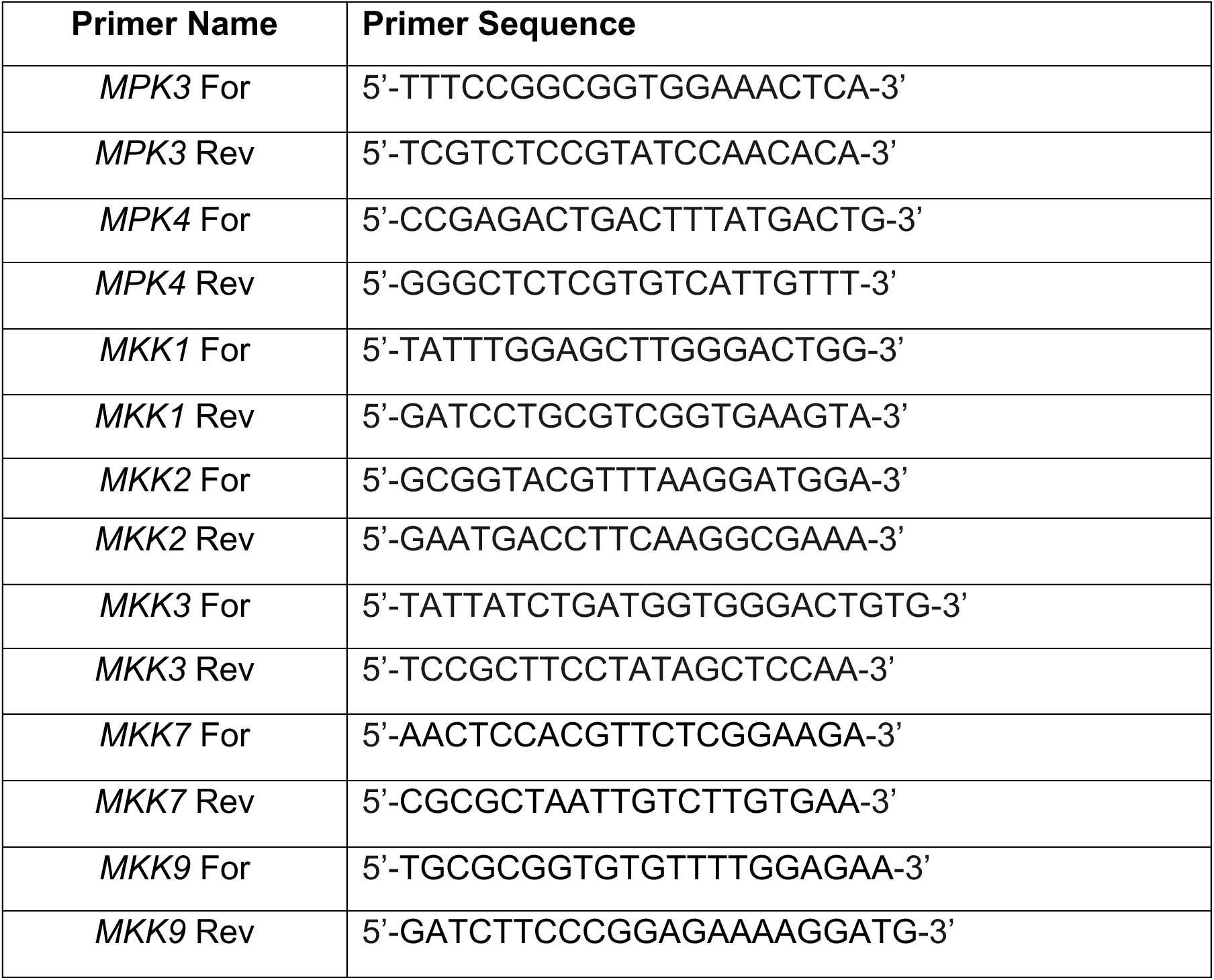
Primer sequences of *MPKs* and *MKKs* used for qRT-PCR.

